# CTLA-4 tail fusion enhances CAR-T anti-tumor immunity

**DOI:** 10.1101/2023.03.14.532655

**Authors:** Xiaoyu Zhou, Hanbing Cao, Shao-Yu Fang, Ryan D. Chow, Kaiyuan Tang, Medha Majety, Meizhu Bai, Matthew B. Dong, Paul A. Renauer, Xingbo Shang, Kazushi Suzuki, Andre Levchenko, Sidi Chen

**Author notes:** Correspondence: SC, (lab). Co-first authors (Xiaoyu Zhou, Hanbing Cao).

## Abstract

Chimeric antigen receptor (CAR) T cells are powerful therapeutics; however, their efficacy is often hindered by critical hurdles. Here, utilizing the endocytic feature of the cytotoxic T-lymphocyte-associated antigen-4 (CTLA-4) cytoplasmic tail (CT), we reprogram CAR function and substantially enhance CAR-T efficacy *in vivo*. CAR-T cells with monomeric, duplex, or triplex CTLA-4 CTs (CCTs) fused to the C-terminus of CAR exhibit a progressive increase in cytotoxicity under repeated stimulation, accompanied by reduced activation and production of pro-inflammatory cytokines. Further characterization reveals that CARs with increasing CCT fusion show a progressively lower surface expression, regulated by their constant endocytosis, recycling and degradation under steady state. The molecular dynamics of reengineered CAR with CCT fusion results in reduced CAR-mediated trogocytosis, loss of tumor antigen, and improved CAR-T survival. CARs with either monomeric (CAR-1CCT) or duplex CCTs (CAR-2CCT) have superior anti-tumor efficacy in a relapsed leukemia model. Single-cell RNA sequencing and flow cytometry analysis reveal that CAR-2CCT cells retain a stronger central memory phenotype and exhibit increased persistence. These findings illuminate a unique strategy for engineering therapeutic T cells and improving CAR-T function through synthetic CCT fusion, which is orthogonal to other cell engineering techniques.

## Introduction

Adoptive cellular immunotherapy with engineered chimeric antigen receptor (CAR) T cells has led to durable clinical responses in hematologic malignancies ^1^. Despite the high response rates, relapse occurs in a large proportion of patients due to a number of factors, including tumor antigen loss, T cell exhaustion, T cell dysfunction, and poor *in vivo* persistence of CAR-T cells ^2, 3^. These are among the series of major challenges with CAR-T therapy, urging for better designs of CARs to overcome the therapeutic hurdles. Currently, the typical structure of a CAR is composed of: (i) a single-chain variable fragment (scFV) for target recognition; (ii) a hinge for flexibility to access the target antigen; (iii) a transmembrane domain for cell surface anchoring; (iv) an intracellular domain containing costimulatory signaling domain(s) (which vary between the 1^st^, 2^nd^ and 3^rd^ generation of CARs); (v) and a T-cell receptor signaling domain (CD3zeta) for signal transduction ^4^. Compared with the 1^st^ generation CAR, better clinical performances have been achieved in subsequent generations by incorporating CD28 ^5^ or 4-1BB ^6^ costimulatory signaling domains. Additionally, weakening the signaling strength of CAR through transient cessation of CAR^7^ or the adoption of low-affinity CAR^8^ has been shown to improve the efficacy of CAR-T cells. These studies highlighted that optimal CAR design is crucial to achieve durable anti-tumor efficacy of CAR-T therapy ^9, 10^.

Trogocytosis is a process by which lymphocytes such as T^11^, B^12^ and natural killer (NK) ^13^ cells extract surface molecules from antigen presenting cells (APCs) through the immune synapse, and present those molecules on their own surface. Consequences of trogocytosis are context dependent. It has been reported that trogocytosis promotes the selection of high affinity cytotoxic T lymphocytes (CTL) by removing major histocompatibility complex (MHC)-peptide complexes from APCs ^14^. However, it also leads to the elimination of reactive CTLs though fratricide when high amounts of MHC-peptide complexes are acquired and presented on their own surface ^11^. Recently, CAR-T cells have also been demonstrated to undergo trogocytosis, which causes tumor antigen loss and fratricide among themselves, compromising their therapeutic efficacy against cancer ^15–17^. It is therefore critical to develop new approaches to overcome CAR mediated trogocytosis for durable CAR-T therapeutic effect. Studies have shown that the combined targeting of tumor antigens ^15^ or the use of a low-affinity CAR ^16^ are able to rescue CAR-T anti-tumor responses from trogocytosis-induced immune escape. However, these approaches are not adaptable or flexible, as they require generating entirely new scFVs for each cancer type.

As an immune checkpoint molecule, cytotoxic T-lymphocyte-associated antigen-4 (CTLA-4) is crucial for maintaining self-tolerance and homeostasis ^18^. Mechanistically, it was initially reported that the immune inhibitory signal of CTLA-4 was transduced through its cytoplasmic tail (CCT), namely by cell-intrinsic inhibition ^19^. However, accumulating evidence indicates that the inhibitory function of CTLA-4 is instead achieved through a cell-extrinsic process termed trans-endocytosis. In trans-endocytosis, CTLA-4^+^ T cells remove and subsequently internalize costimulatory molecules CD80/CD86 from APCs, rather than presenting them on the cell surface (as in trogocytosis). As a result, activation of other naïve T cells through CD28 signaling is inhibited ^20–23^. To mediate this process, a key feature of CTLA-4 is that it is highly endocytic, such that the majority of CTLA-4 is constantly cycling between the cell surface and the intracellular compartment. This cycling occurs through the interaction between the YVKM motif in CCT and clathrin adaptor activating protein 2 (AP-2), resulting in limited surface expression of CTLA-4. Thus, rather than directly transducing suppressive signals, CCT is believed to regulate the surface availability of CTLA-4 for optimal T cell activation ^22, 24^.

We then wondered if CCT could be utilized for optimal design of CAR with low trogocytosis, thereby improving anti-tumor efficacy of CAR-T therapy. Fusing synthetic CCTs to the C-terminus of the CAR led to progressively decreased surface CAR expression, mediated by ongoing endocytosis, recycling and degradation under steady state. CAR-CCT cells had reduced trogocytosis while further demonstrating enhanced cytolytic activity against cognate cancer cells *in vitro,* with decreased activation and production of proinflammatory cytokines. Compared to control CAR-T cells, CAR-CCT-T cells have significantly reduced tonic signaling and increased responsiveness to antigen stimulation at transcriptome level. In addition, CAR-T cells with either monomeric (CAR-1CCT) or duplex CCTs (CAR-2CCT) have more potent and durable anti-tumor efficacy *in vivo*. Further *in vivo* immune characterization showed that CAR-2CCT-T cells are more persistent and exhibit enhanced central memory differentiation. Together, these data illuminate a distinct approach to engineer CAR-T cells via synthetic fusion of CCTs to CARs. Synthetic CCT fusion confers enhanced CAR function independently of the specific CAR type and is broadly compatible with other CAR engineering approaches.

## Results

### Engineered CAR with CCT fusion confers CAR-T cell durability under repeated stimulations

To test whether CCT engineering can improve the functionality of CAR-T cells, we designed constructs to express a CD22-CAR (hereafter abbreviated CAR) based on the m971-BBz backbone, with monomeric (CAR-1CCT), duplex (CAR-2CCT), or triplex CCTs (CAR-3CCT) fused to the C terminus (**Fig. 1a**). In all of these CAR constructs, we also encoded a Flag tag at the N terminus for surface CAR detection, along with an mScarlet reporter for monitoring viral transduction. We transduced human CD3 T cells with lentiviruses carrying the above constructs to generate CAR-T cells (**Fig. 1a**). Transduction with different constructs did not affect the CD4/CD8 T cell composition (**Fig. S1a and b**).

**Figure 1.**
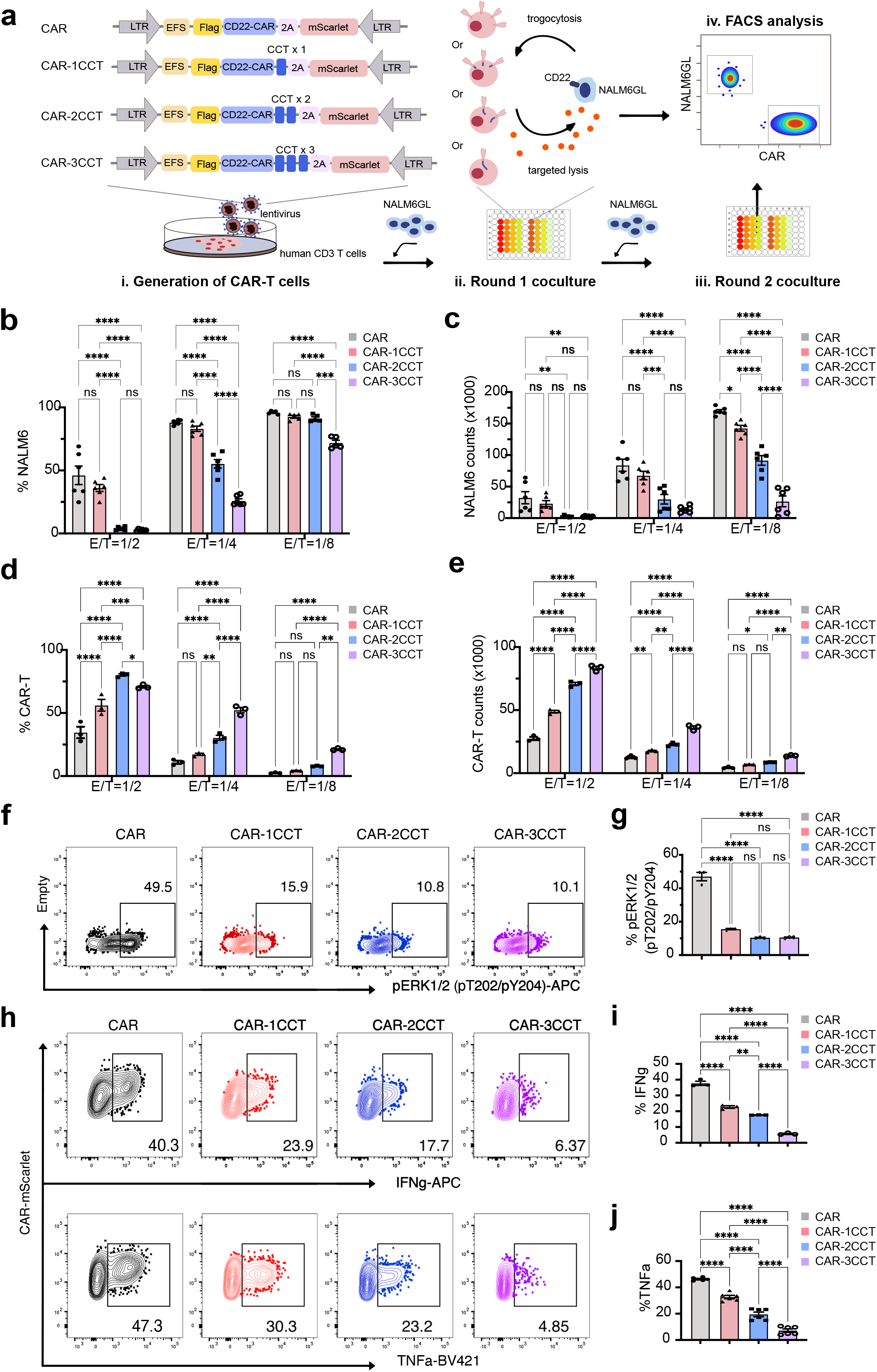
Engineering CAR with CCT fusion increased persistent CAR T cell function under repeated tumor stimulation. **a.** Schematics of the workflow used to generate CAR-T cells for effector function assessment *in vitro*. (i) Generation of CAR-T cells. Human CD3 T cells were infected by lentiviruses carrying CAR targeting human CD22 (m971-CD28-41BB-CD3zeta). These CARs were engineered with either monomeric (CAR-1CCT), duplex (CAR-2CCT) or triplex CCTs (CAR-3CCT) fused to the C-terminal of human CD22 CAR (CAR), followed by a fluorescent reporter mScarlet, separated by T2A sequences. Flag sequences were tagged at N-terminal of the CAR for surface expression detection. (ii) Round 1 coculture. Those four groups of CAR-T cells generated in step (i) were coculture with NALM6GL cells at variable E/T ratios for 24 hours. (iii) Round 2 coculture. An additional round of NALM6GL cells were supplemented into the the initial coculture for 24 hours. (iv). FACS analysis. Cells from both round1 and 2 cocultures were quantified by fluorescence-activated single cell sorting (FACS). **b.** Quantification of NALM6GL percentage after 2 rounds of NALM6GL stimulation. **c.** Quantification of NALM6GL counts after 2 rounds of NALM6GL stimulation. **d.** Quantification of CAR-T cell percentage after 2 rounds of NALM6GL stimulation. **e.** Quantification of CAR-T cell counts after 2 rounds of NALM6GL stimulation. **f.** Representative FACS plots demonstrating the expression of pERK1/2(pT202/pY204) after crosslinking CAR-T cells with CD22-biotin and streptavidin at 37°C for 5 minutes. All CAR-T cells were previously gated on mScarlet^+^ populations for quantification. **g.** Quantification of pERK1/2(pT202/pY204) expression in CAR-T cells as shown in (f). **h.** Representative FACS plots demonstrating the intracellular expression of IFNg and TNFa by CAR-T cells after stimulated with NALM6GL cells at an E/T ratio of 1:1 for 4 hours. **i and j**, Quantification of intracellular IFNg (**i**) and TNFa (**j**) expression as shown in **(h)**. For all bar plots, data are shown as mean ± s.e.m. One-way ANOVA with Dunnett’s multiple-comparisons test is used to assess significance for figure (**i**) and (**j**). Two-way ANOVA with Tukey’s multiple-comparisons test is used to assess significance for figure (**b-e**). ns = *p* > 0.05, * = *p* < 0.05, ** = *p* < 0.01, *** = *p* < 0.001, **** = *p* < 0.0001.

We assessed the effector function of the engineered CAR-CCT cells by coculturing them with NALM6 cells (B cell leukemia) expressing GFP-luciferase (NALM6GL), followed by simultaneous quantification of NALM6GL and CAR-T cells using flow cytometry (**Fig. S1c**). All four groups of CAR-T cells lysed approximately 100% of NALM6GL when cocultured with one round of cancer cells for 24 hours (**Fig. S1d**). However, this was achieved with distinct consequences, with significantly more CAR-T cells surviving when expressing a CAR-CCT fusion construct (CAR-1CCT, CAR-2CCT and CAR-3CCT) (**Fig. S1e**). Accordingly, we observed significantly lower expression of T cell exhaustion markers LAG-3 and PD-1 in CAR-2CCT and CAR-3CCT cells compared to the control CAR-T cells (**Fig. S1f-i**). Other coinhibitory markers such as TIGIT, cycling CTLA-4, and recycling CTLA-4 were comparable between groups; TIM3 was undetectable in all groups (**Fig. S1j-o**).

We then hypothesized that engineered CAR with CCT fusion might support the persistent anti-tumor response of T cells under repeated antigen stimulation, which is commonly seen in tumor relapses. To test this hypothesis, a second round of NALM6GL was introduced 24 hours after the initial coculture (**Fig. 1a**). Compared with the control CAR-T cells, CCT fusion in all three groups resulted in markedly fewer live NALM6GL cells, quantified as either percentage (**Fig. 1b and S2a)** or counts (**Fig. 1c and S2a**), indicating the superior capability of CAR-CCT cells to kill NALM6GL cells. Similar to what had been observed after the first round of NALM6GL stimulation, significantly more CAR-T cells in the CAR-CCT groups survived the second round NALM6GL stimulation (**Fig. 1d and e**), with the same number of input CAR-T cells seeded at the beginning of the co-culture (**Fig. S2b**). Notably, after two rounds of cancer stimulation, CAR- 2CCT and CAR-3CCT cells showed higher expression of central memory markers such as CD45RO and CD62L (**Fig. S2c and S2d**), while maintaining lower LAG-3 expression (**Fig. S2e and S2f**) than the control CAR-T cells.

To investigate the impact of tumor antigen density on this phenotype seen with CAR-CCT cells, we generated NALM6-CD22-GFP cells, in which CD22 was tagged with GFP by a GGGGS linker. CD22^high^ and CD22^low^ NALM6 cells were sorted out based on the CD22 expression level. *In vitro* coculture of CAR-CCT cells with either CD22^high^ or CD22^low^ NALM6 cells showed that CAR-CCT cells induced more efficient killing than the control CAR-T cells, with CAR-2CCT and CAR-3CCT showing the highest capability of killing both CD22^high^ and CD22^low^ NALM6 cells (**Fig. S2g-i**). When repeatedly challenged with NALM6 cells, all four groups of CAR-T cells were more effective at killing cancer cells with lower CD22 expression (**Fig. S2g-i**). These data suggest that, at least in the context of an *in vitro* co-culture system, a high cancer antigen load is unfavorable for CAR-T function under repeated challenge. Altogether, these results suggest that CAR-T cells with engineered CCT fusion have improved durability under repeated antigen stimulations, with enhanced cytolytic ability and a stronger central memory phenotype.

As our approach involves engineering the distal cytoplasmic end of the CAR, we sought to confirm its versatility by engineering CCT fusion to CAR targeting a different antigen. By replacing the m971 scFv with the FMC63 scFv, we similarly engineered a series of CAR and CAR-CCT constructs with a human CD19 targeting scFv (FMC63), namely CD19 CAR (19CAR), as well as monomeric (19CAR-1CCT), duplex (19CAR-2CCT), and triplex CCTs (19CAR-3CCT) (**Fig. S3a**). We repeated the experiments above with this series of 19CARs and observed similar results, where significantly more 19CAR-T cells survived the two rounds NALM6GL coculture (**Fig. S3b and 3c**), which progressively killed more NALM6GL cells with increasing number of CCTs (**Fig. S3b and 3d**). Collectively, these results demonstrated that harnessing CCT fusion to enhance CAR-T function is applicable to more than one type of CAR.

### CAR-T cells with CCT fusion show decreased activation and production of pro-inflammatory cytokines

We additionally investigated if CCT fusion altered CAR signaling by checking ERK1/2 phosphorylation and cytokine production after stimulating CAR-CCT cells with tumor antigen. We found that all CAR-CCT were responsive to antigen stimulation, and increasing CCT fusion decreased the expression level of phospho-ERK1/2 (pT202/pY204), IFNg and TNFa (**Fig. 1f-j**). We also examined the secretion of other cytokine and cytotoxic molecules. We quantified the secreted proteins from the supernatants of NALM6 cocultures with the control CAR or CAR-CCT cells by LEGENDPlex bead-based immunoassays. Our results showed that CCT-engineered CAR- T cells, especially the CAR-3CCT group, had elevated levels of granzymes, perforin and granulysin (**Fig. S3e**), which are the main factors for targeted killing by CAR-T cells^25^. On the other hand, CCT-engineered CAR-T cells, in all three groups, had reduced levels of pro-inflammatory cytokines, such as IL-4, IL-6, IL-2, TNF-a, IL17a and IFNg (**Fig. S3e**), all of which are associated with cytokine release syndrome seen in clinical CAR-T therapy ^26, 27^. These data suggest that engineered CAR-CCT cells have improved killing capability with elevated degranulation, but decreased ERK phosphorylation, IFNg and TNFa production. Given the increased *in vitro* killing ability and elevated granzymes, CAR-CCT cells are more efficient in their ability to kill cancer cells. The accompanying decrease in ERK phosphorylation, IFNg, and TNFa production may imply a reduction in T cell activation and inflammatory responses, possibly due to alterations in CAR dynamics when CCT is fused. The phenomenon that engineered T cells with enhanced cytotoxicity but low cytokine release have also been reported by others ^28^, potentially limiting the risk of cytokine release syndrome and cerebral edema/neurotoxicity, two of the major side effects associated with current CAR-T cell therapy ^29^.

### CCT fusion enables titration of CAR expression through receptor endocytosis, recycling and degradation at steady state

CTLA-4 is constantly internalized through endocytosis, after which it is either recycled back to the cell surface or degraded ^30–32^. To gain insight into the improved killing ability and reduced activation of CAR-CCT cells, we sought to determine if CARs with CCT fusion also possess similar molecular dynamics as native CTLA-4. We first observed that CCT fusion reduced surface CAR expression in a dose-dependent manner (number of fused CCTs), as indicated by Flag staining after gating on mScarlet^+^ CAR-T cells (**Fig. 2a and 2b**). This reduction in surface CAR expression may explain the decreased activation level seen in CAR-CCT cells following antigen stimulation (**Fig. 1f-j**). To further characterize the cellular localization pattern of CAR-CCT molecules, we performed a flow cytometry staining procedure that distinguishes between endocytic and surface CAR (**Fig. 2c**). CARs with CCT fusion had altered cellular localization patterns and displayed endocytic features, with CAR-1CCT displayed the highest endocytosis level (**Fig. 2d and 2e)**. Analogous to the results in **Fig. 2a**, we again observed that with increasing CCT fusion, surface CAR levels decreased. CAR-3CCT cells had the highest number of mScarlet^+^ CAR^-^ populations (**Fig. 2d and 2e)**, suggestive of elevated degradation of CAR-3CCT molecules.

**Figure 2.**
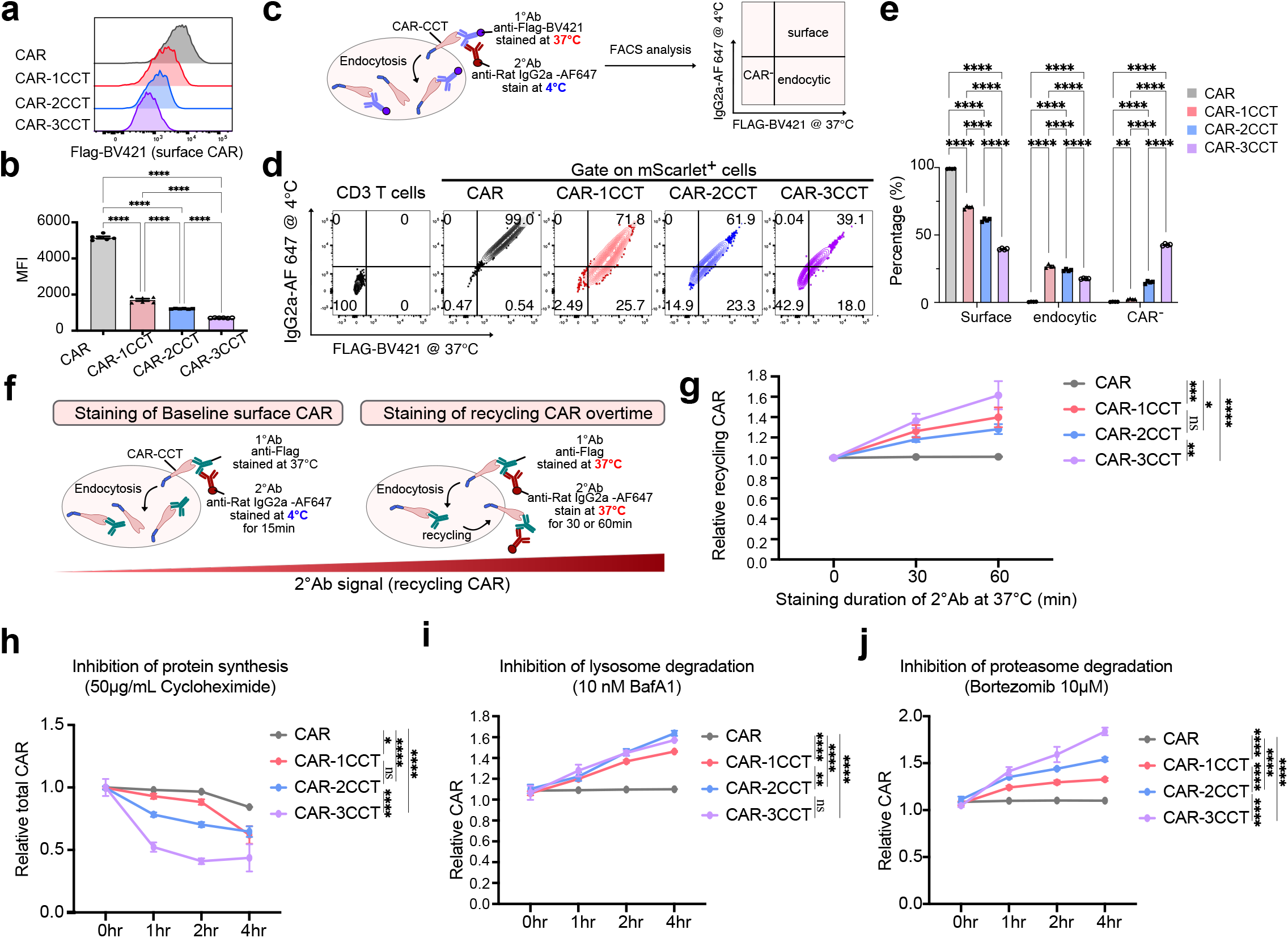
CCT fusion enables titration of CAR expression through receptor endocytosis, recycling and degradation at steady state. **a.** Representative flow cytometry analysis of CAR-T cells, pre-gated on mScarlet^+^ populations, showing surface expression level of CAR, indicated by Flag staining. **b.** Quantification of **a** showing surface flag expression in medium fluorescent intensity (MFI). **c.** Schematics of the workflow used in **d** and **e** for detecting CAR endocytosis. Surface and endocytic CAR were first stained with primary anti-Flag-BV421 at 37°C for 30 minutes, followed by a secondary anti-Rat IgG2a-AF647 stained at 4°C for 15 minutes. The surface, endocytic CAR and CAR^-^ populations were differentiated based on their staining patterns of the primary and secondary antibodies. **d.** Representative flow cytometry analysis of CAR-T endocytosis. **e.** Quantification of **d** showing the level of CAR-T endocytosis. **f.** Schematics of the workflow used in **g** for detecting recycling CAR. Surface and endocytic CAR were first stained with primary anti-Flag at 37°C for 30 minutes, followed by a secondary anti-Rat IgG2a-AF647, either stained at 4°C for 15 minutes (baseline) or at 37°C for 30 minutes or 60 minutes. Recycling was quantified by the increase in the percentage of cells stained positive for the secondary antibody. **g.** Quantification of recycling CAR over time. **h.** Quantification of the stability CAR with the presence of 50µg/mL Cycloheximide by intracellular staining of Flag. Relative CAR expression was determined by dividing the proportion of Flag^+^ CAR-T cells after treatment using Cycloheximide with those treated using DMSO (0 hr). **i-j**. Quantification of CAR expression in the presence of 10nM Bafilomycin A1 (BafA1) (**i**) or 10µM bortezomib (**j**) by performing intracellular staining of CAR-T cells that had been previously fed with anti-flag antibody at 37°C for 1 hour, followed by detection using anti-Rat-IgG2a- Alexaflour 647 specific to the primary antibody. Relative CAR expression was determined by dividing the proportion of Alexaflour 647^+^ CAR-T cells after treatment using degradation inhibitors with those treated using DMSO (0 hr). For all bar plots, data are shown as mean ± s.e.m. One-way ANOVA with Dunnett’s multiple-comparisons test is used to assess significance for figure (**b**). Two-way ANOVA with Tukey’s multiple-comparisons test is used to assess significance for figure (**e**) and (**g-j**). ns = *p*> 0.05, * = *p* < 0.05, ** = *p* < 0.01, *** = *p* < 0.001, **** = *p* < 0.0001.

To characterize the recycling pattern of the CAR-CCT molecules, we further modified the staining protocol. Specifically, we compared the levels of staining signal when the secondary antibody was incubated at 4°C for 15 minutes or at 37°C for either 30 or 60 minutes (**Fig. 2f**). Compared to the 4°C incubation, an increase of staining at 37°C for either 30 or 60 minutes is indicative of active recycling CAR molecules. We found that in contrast to the control CAR, all CAR-CCT molecules showed recycling features, with CAR-3CCT showing the highest recycling rate, and CAR-1CCT and CAR-2CCT demonstrating similar recycling rates (**Fig. 2g**).

We additionally quantified the stability of the CAR-CCT molecules at steady state. By inhibiting de novo protein translation with cycloheximide ^33^, we found that CAR-CCT molecules had significantly decreased stability, with CAR-3CCT molecules demonstrating the highest degradation rate (**Fig. 2h**). Next, we sought to investigate the pathway through which CAR-CCT molecules were being degraded. As native CTLA-4 is subjected to both lysosomal and proteasome degradation ^34^, we investigated their contribution to CAR-CCT stability by introducing Bafilomycin A1(BafA1, lysosomal inhibitor) or bortezomib (proteasomal inhibitor). BafA1 or bortezomib treatment significantly increased relative CAR expression in CAR-CCT groups over time, while no obvious change was observed in the control CAR group, suggesting that both the lysosome and proteasome are involved in CAR-CCT degradation (**Fig. 2i-j**). Interestingly, we observed a CCT fusion dose-dependent effect when the cells were treated with bortezomib, indicating that with increasing CCT fusions, proteasomal degradation of the CAR-CCT is also increased (**Fig. 2j**). In contrast, there was no difference in CAR expression between CAR-3CCT and CAR-2CCT when lysosomal degradation was inhibited by BafA1 (**Fig. 2i**), though 2CCT and 3CCT were nevertheless subject to more lysosomal degradation than 1CCT or the baseline CAR. These results suggest that CCT fusions promote lysosomal degradation, but only up to certain point (2CCT); on the other hand, increasing CCT fusions progressively promote proteasomal degradation at least up to 3CCTs. Collectively, these data suggest that the surface expression of CAR-CCT molecules is regulated by their constant endocytosis, recycling and degradation under steady state, contrasting with that of the control.

### CCT fusion reduces CAR-mediated trogocytosis

CAR-T cells have been shown to acquire surface molecules from tumor cells through trogocytosis, leading to antigen loss that compromises anti-tumor efficacy ^15^. In line with these prior observations, we performed time-lapse live cell imaging and found that CAR-T cells actively acquired CD22 antigen from NALM6GL cells (**Supplemental video 1**). By including a dye that only penetrates dead cells, we recorded active fratricide among CAR-T cells, as indicated by influx of the dye after an active engagement between CD22^+^ CAR-T cells (**Fig. 3a and Supplemental video 1**). Moreover, we found that latrunculin A, an F-actin inhibitor^15, 35^, inhibit the transfer of CD22 antigen from NALM6 cells to T cells, a key hallmark of CAR-mediated trogocytosis (**Fig. S4a and S4b**), which further support the observations.

**Figure 3.**
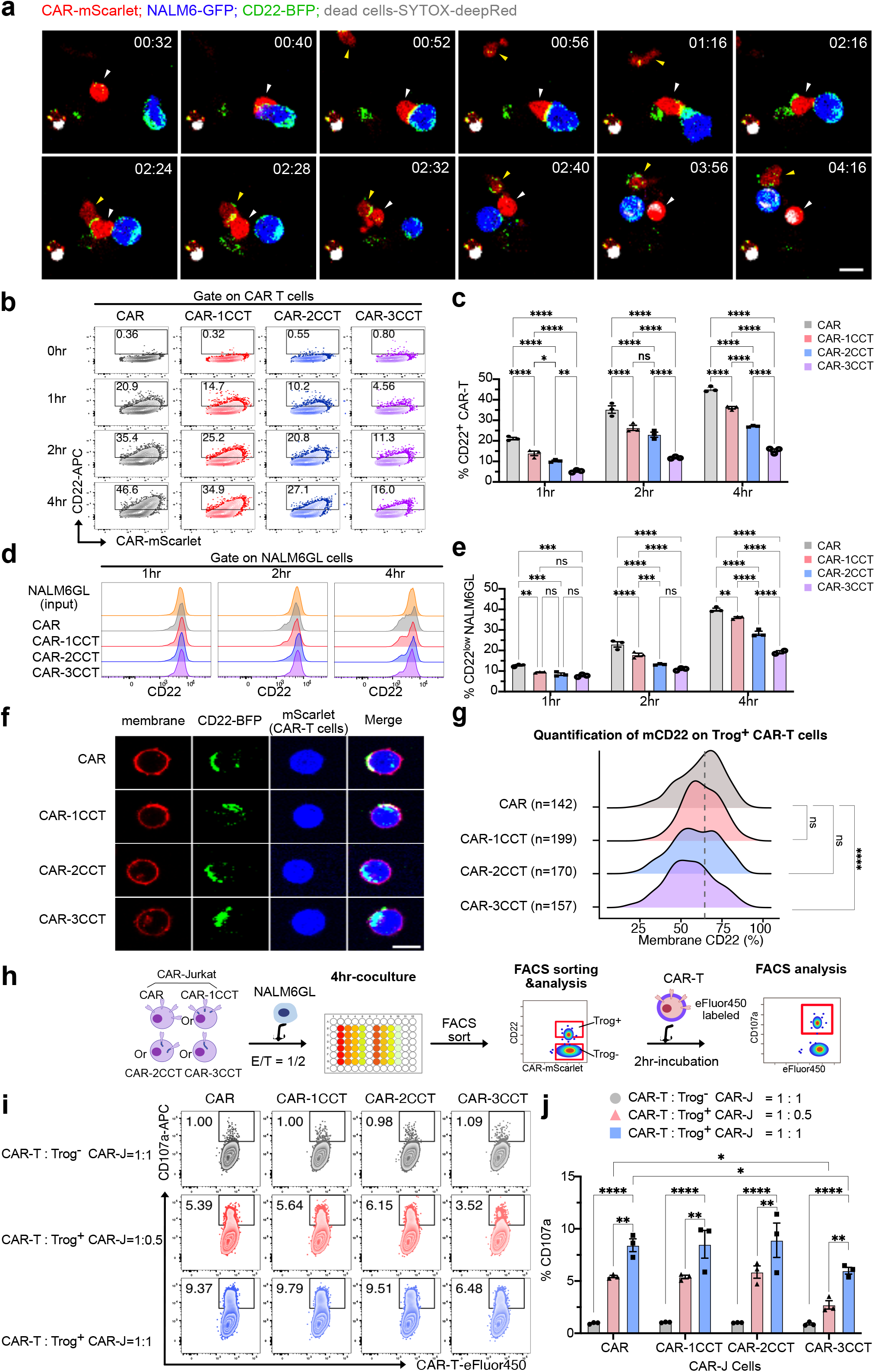
CCT fusion effectively decreases CAR-mediated trogocytosis and accompanying fratricide. **a.** Representative time-lapse live cell images showing trogocytosis and fratricide of CAR-T cells after introduction with NALM6GL-CD22-BFP cancer cells. The white arrow labels a CAR-T cell that acquired CD22-BFP after an active engagement with a cancer cell, which subsequently rendered it susceptible to CAR-T fratricide afterwards, as indicated by the influx of SYTOX deep red dye. This white arrow-labeled cell then contacted another CAR-T cell (yellow arrow), interacting at a region with enhanced CD22-BFP signal. Shortly after this active interaction, the white arrow-labeled cell was stained by the SYTOX deep red dye, indicating cell death. The time stamps on the top right of each image are represented as hour: minute, scale bar = 10µm. **b.** Representative flow cytometry results showing CD22 expression on CAR-T cells after 0-4 hours coculturing with NALM6GL cells at E/T ratio = 1/1. **c.** Quantification of **b** showing CD22 expression on CAR-T cells. **d.** Representative flow cytometry results showing CD22 expression on NALM6GL cells after 0-4 hours coculturing with CAR-T cells at E/T ratio = 1/1. **e.** Quantification of **d** showing CD22 expression on NALM6GL cells. **f.** Representative confocal images showing the CD22-BFP colocalization with surface membrane of Trog^+^ CAR-T cells. Scale bar = 10µm. **g.** Quantification of **f** showing the distribution of % membrane CD22 (mCD22). In each group, the total number of quantified cells is labelled on the Y-axis of the plot. Dash line represents the median percentage of mCD22 on the control CAR-T cells. **h.** Schematics of the workflow used in **i** and **j** for detecting the degranulation of CAR-T cells. CAR-Jurkat (CAR-J) cells were cocultured with NALM6-CD22-GFP at an E/T ratio of 1:1 for 4 hours to allow for CD22-GFP transfer from NALM6 to CAR-Jurkat cells. Both Trog^+^ CAR-J and Trog^-^ CAR-J cells were sorted out based on their expression of CD22-GFP. Those cells were then cocultured with eFluor450 labelled CAR-T cells for 2 hours for determine the expression of CD107a. **i.** Representative flow cytometry results showing the CD107a expression of fresh CAR-T cells in coculture with Trog^+^ or Trog^-^ CAR-J cells at various E/T ratios. **j.** Quantification of **i**. For all bar plots, data are shown as mean ± s.e.m. One-way ANOVA with Dunnett’s multiple-comparisons test is used to assess significance for figure (**g**). Two-way ANOVA with Tukey’s multiple-comparisons test is used to assess significance for figure (**c**), (**e**) and (**j**). ns = *p*> 0.05, * = *p* < 0.05, ** = *p* < 0.01, *** = *p* < 0.001, **** = *p* < 0.0001.

As it has been reported that trogocytosis is impacted by CAR affinity^36^, we wondered whether the altered molecular dynamics of CAR-CCT might also impact its propensity for mediating trogocytosis. From *in vitro* cocultures of NALM6GL and CAR-T cells, we detected substantial surface CD22 antigen on CAR-T cells that normally do not express CD22, as early as 1 hour after incubation (**Fig. 3b and 3c**), consistent with rapid trogocytosis by CAR-T cells. Engineered CCT(s) in CARs significantly reduced trogocytosis-mediated acquisition of surface CD22 in a CCT-dose-dependent fashion (**Fig. 3b-c, S4a and S4b**). Concordantly, the NALM6GL cancer cells retained progressively more surface CD22 antigen when co-cultured with CAR-T cells harboring increasing numbers of engineered CCTs (**Fig 3d and e**). These results showed that CCT fusion efficiently reduces CAR-mediated trogocytosis and CD22 loss in cancer cells.

As CTLA-4 has been shown to internalize ligands acquired by trans-endocytosis ^21^, we next quantified the localization of the transferred CD22 antigen by sorting out CD22^+^ CAR-T (Trog^+^) cells for confocal imaging. In all groups, the majority of the transferred CD22 antigen localized to the cell membrane (**Fig. 3f and 3g**). Of note, only very high levels of CCT fusion (3CCT) seemed to decrease membrane colocalization, given that a significant leftward shift of the membrane CD22(%) distribution was only observed for CAR-3CCT (**Fig. 3g**). Thus, the presence of 1CCT or 2CCT fusion does not significantly change the surface localization of CD22 molecules obtained through CAR-mediated trogocytosis, whereas CAR-3CCT fusion leads to increased internalization.

To determine if the CD22 antigen acquired by trogocytosis is still functional in stimulating CAR- T cells, we generated CAR-Jurkat (CAR-J) cells, which undergo trogocytosis of target CD22 (**Fig. S4c and S4d**) without killing the target cells ^37^. After transducing Jurkat cells with different CAR- CCT constructs, we incubated them with NALM6-CD22-GFP cells to allow for CD22-GFP transfer onto Jurkat cells. Both CD22-GFP^+^ (Trog^+^) and CD22-GFP^-^ (Trog^-^) CAR-J cells were sorted out and used as target cells to stimulate fresh CAR-T cells (effector cells) that had been labeled with eFluor450 (**Fig. 3h**). In all groups, Trog^+^ CAR-J cells could induce the cytotoxic degranulation of CAR-T cells, as indicated by the membrane expression of CD107a. This was achieved in a dose-dependent manner, as higher degranulation occurred when more target Trog^+^ CAR-J cells were added to the co-culture (**Fig. 3i and 3j**).

Given that significantly lower percentages of Trog^+^ were observed in the CAR-CCT-T cells after their encounter with cancer cells (**Fig. 3b and 3c**), lower degranulation, and thus lower fratricide among them could be inferred. In comparison, Trog^-^ CAR-J cells did not meaningfully induce degranulation, as expected (**Fig. 3j**). These data indicate that CD22 transferred through trogocytosis is functional in stimulating CAR-T cell degranulation. While the stimulation of CAR- T degranulation was similar among Trog^+^ control CAR-J cells, CAR-1CCT-J and CAR-2CCT-J cells, Trog^+^ CAR-J cells with 3CCT fusion induced significantly lower degranulation (**Fig. 3i and 3j**). This may result from the enhanced internalization of transferred CD22 by CAR-3CCT, as seen in **Fig 3f and 3g**. Together, these results suggest that CCT fusion quantitively reduces CAR- mediated trogocytosis, loss of cancer antigen.

### CCT fusion enhances the survival and proliferation of CAR-T cells

Given our observation that CAR-CCT cells were more abundant after repeated antigen stimulation and also exhibited reduced trogocytosis, we wondered whether CCT fusion impacted CAR-T cell survival and proliferation. To test this, we mixed CAR-CCT cells with control CAR-T cells labeled with eFluor450 dye, followed by NALM6GL stimulation. This ensured that both the CAR-CCT and control CAR-T cells were subjected to the same stimulation (**Fig. 4a**). Quantifying the relative abundance of eFluor450^high^ and eFluor450^low^ cells, we found that CAR-T cells with CCT fusion exhibited improved survival than the control (**Fig. 4b-d**). To assess the potential role of cell proliferation in these findings, we used flow cytometry and eFluor450 labeling to quantify CAR- T cell proliferation following stimulation with NALM6GL cells. CAR-2CCT and CAR-3CCT cells exhibited greater proliferation potential, as more cells underwent 4 or 5 rounds of division in these two groups (**Fig. 4e and 4f**). In an analogous co-culture experiment, we further observed that Trog^+^ CAR-T cells were significantly more apoptotic than the Trog^-^ CAR-T cells, and CAR-T cells with increasing CCT fusion had reduced apoptosis levels (**Fig. 4g-i**). These results suggest that the high levels of trogocytosis induced by the control CAR are unfavorable for T cell survival, and that CAR-CCT fusion inhibits this process to improve CAR-T survival following antigen exposure.

**Figure 4.**
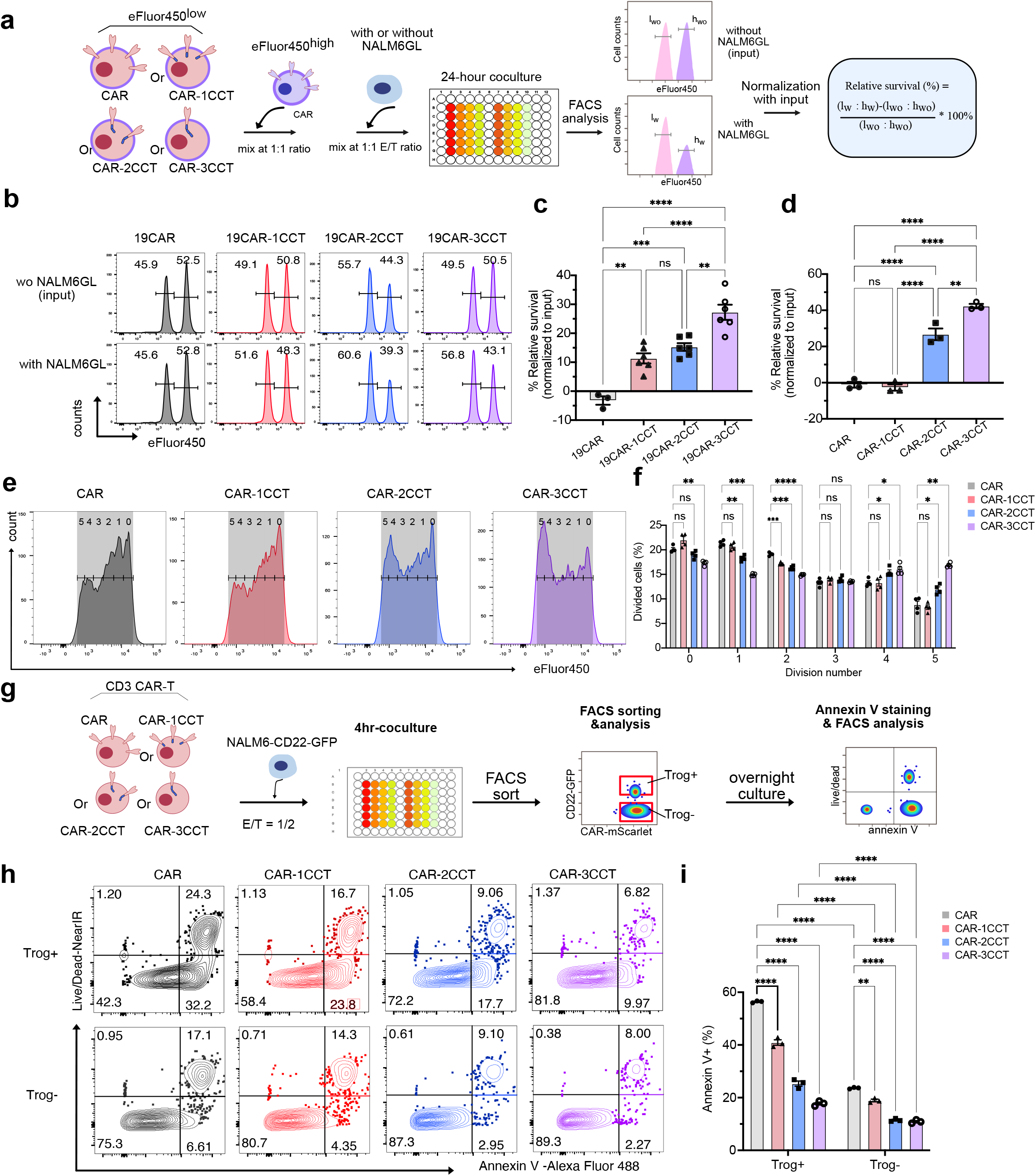
CAR-T cells with CCT fusion show increased survival and proliferation *in vitro* upon stimulation with cancer cells. **a.** Schematics of the CAR-T survival assay used in **b-d**. Control CAR-T cells labeled with 10µM eFluor450(eFluor450^high^), were mixed 1:1 with CAR, CAR-1CCT, CAR-2CCT or CAR-3CCT cells labeled with 1µM eFluor450(eFluor450^low^). These mixed CAR T cells were then cocultured with or without NALM6GL cells at E/T = 1/2 for 24 hours. Relative percentages of eFluor450 high and low populations gated from CAR^+^ (mScarlet^+^) cells were used for quantification of relative survival of CAR-T cells. Relative CAR-T survival (%) = [(lw : hw)-(lwo : hwo)]/ (lwo : hwo)*100% , in which lw and hw stands for percentage of eFluor450 low and high populations, respectively, with NALM6GL cells stimulation, while lwo and hwo stands for percentage of eFluor450 low and high populations, respectively, without NALM6GL cells stimulation. **b.** Representative flow cytometry results showing the relative survival of 19CAR-T cells from the assay described above. **c.** Quantification of **b** showing the relative survival of 19CAR-T cells. **d.** Quantification of relative survival of CAR-T cells (CD22-CAR-T) from the assay described above. **e.** Representative flow cytometry results showing the dilution of proliferation staining dye (eFluor450). Purified CAR-T cells were stained with 10 µM eFluor450 and cultured in cX-vivo medium for 4 days before analyzed by flow cytometry. The number labeled on the plot indicates the division number modulated using FlowJo. **f.** Quantification of **e** showing the percentage of proliferated CAR-T cells corresponding to each division number. **g.** Schematics of the workflow used in **h-i**. CAR-T cells were cocultured with NALM6-CD22- GFP cells at E/T = 1/2 for 4 hours. Both Trog^+^ CAR-T and Trog^-^ CAR-T cells were sorted out based on their expression of CD22-GFP, and cocultured separately overnight before staining for Annexin V. **h.** Representative flow cytometry results showing the Live/Dead and Annexin V staining of sorted Trog^+^ and Trog^-^ CAR-T cells. **i.** Quantification of **h** showing the percentage of Annexin V^+^ CAR-T cells. For all bar plots, data are shown as mean ± s.e.m. One-way ANOVA with Dunnett’s multiple-comparisons test is used to assess significance for figure (**c**) and (**d**). Two-way ANOVA with Tukey’s multiple- comparisons test is used to assess significance for figure (**f**) and (**i**). ns = *p*> 0.05, * = *p* < 0.05, ** = *p* < 0.01, *** = *p* < 0.001, **** = *p* < 0.0001.

We next investigated whether the enhanced survival of CAR-CCT cells was also evident in the *in vivo* setting. In mice intravenously injected with NALM6GL cells, adoptively transferred CAR-T cells from bone marrow, where most cancer cells reside ^38^, were harvested for analysis. We found that CAR-T cells engineered with CCT showed significantly higher relative survival (**Fig. 5a-c, S4e**) and lower trogocytosis levels (**Fig. 5d**) compared to those without CCT. Interestingly, we observed a strong correlation between the relative survival and trogocytosis of the CAR-T cells (**Fig. 5e**), with higher survival corresponding to lower trogocytosis. This was consistent with our results showing higher CAR-T: NALM6GL ratios in CAR-T cells engineered with 1CCT or 2CCTs compared to the control CAR-T cells (**Fig. 5f**). These findings collectively demonstrate that CCT fusion effectively improves the survival and proliferation of CAR-T, and impairs the high level of apoptosis associated with trogocytosis.

**Figure 5.**
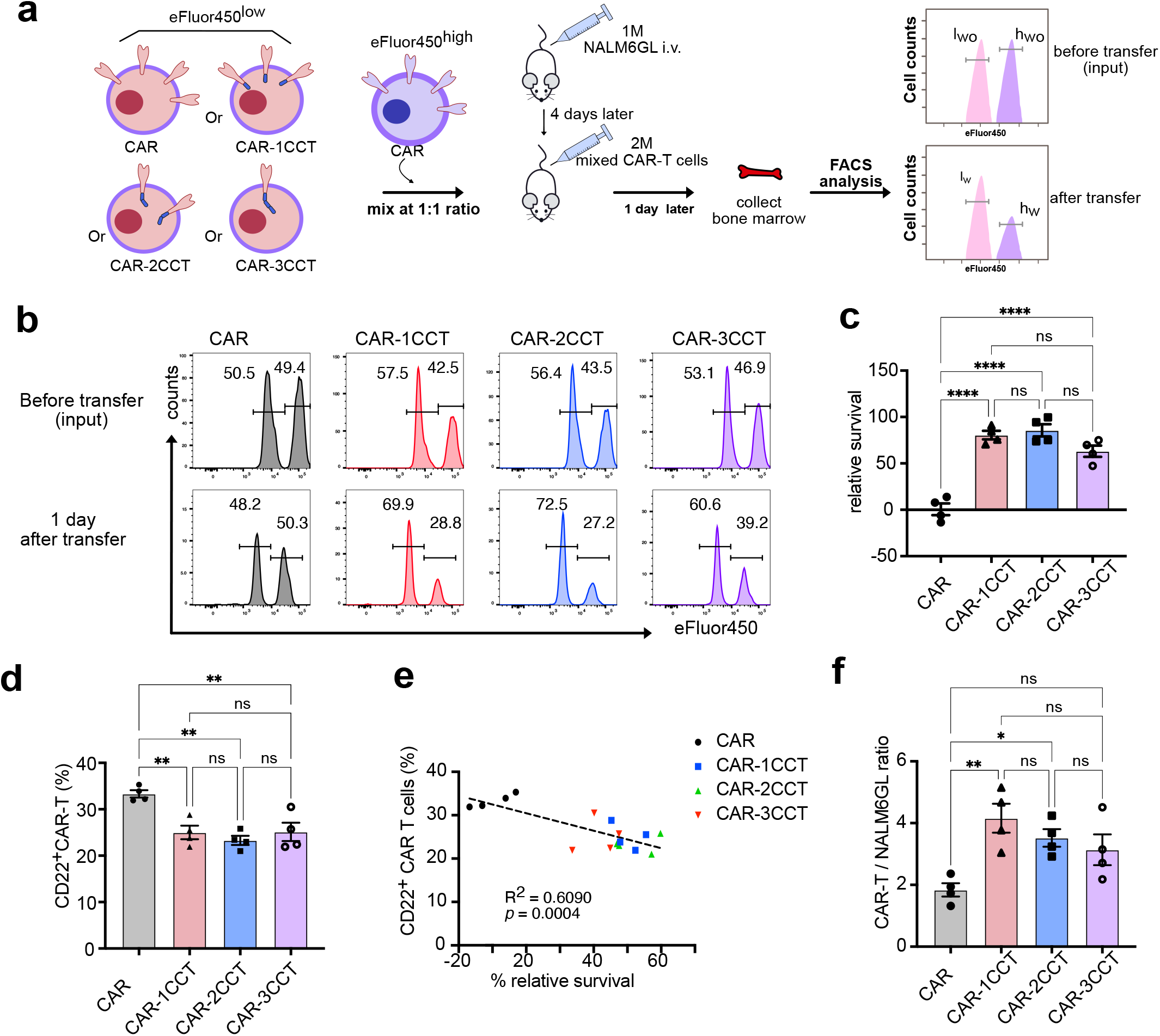
CAR-T cells with CCT fusion show improved survival *in vivo*. **a.** Schematics of *in vivo* experimental workflow used in **b-f**. NSG mice (female, 6-10 weeks old) were first inoculated with 1 million NALM6GL cells intravenously (i.v.) on day 0. CAR-T cells stained with 1µM eFluor450 were mixed with CAR-T cells stained with 10µM eFluor450 at 1:1 ratio. Two million of these labeled cell mixtures were transferred to the leukemia NSG models on day 4. One day post CAR-T transfer, bone marrow samples were collected for flow cytometry analysis to determine the relative percentage of eFluor450^low^ and eFluor450^high^ populations. **b.** Representative flow cytometry results showing the relative survival of CAR-T cells from the assay described above. **c.** Quantification of **b** showing the relative survival of CAR-T cells. Relative survival (%) = [(l_w_ : h_w_)-(l_wo_ : h_wo_)]/ (l_wo_ : h_wo_)*100% , in which l_w_ and h_w_ stands for percentage of eFluor450 low and high populations, respectively, after transfer, while l_wo_ and h_wo_ stands for percentage of eFluor450 low and high populations, respectively, before the transfer. **d.** Quantification of CAR-T trogocytosis *in vivo* as indicated by the CD22 expression detected by staining CAR-T cells from above bone marrow. **e.** Correlation between relative survival and trogocytosis of CAR-T cells. **f.** Quantification of CAR-T cell and NALM6GL relative abundance by calculating the CAR-T: NALM6GL ratio. For all bar plots, data are shown as mean ± s.e.m. One-way ANOVA with Dunnett’s multiple- comparisons test is used to assess significance for figure (**c**), (**d**) and (**f**). ns = *p*> 0.05, * = *p* < 0.05, ** = *p* < 0.01, *** = *p* < 0.001, **** = *p* < 0.0001.

### CAR-T cells with CCT fusion show reduced tonic signaling and increased responsiveness to repeated stimulations at transcriptome level

We characterized CAR-T cells using transcriptome profiling to gain further insight into how CCT fusion engineering impacts their function. We performed mRNA-seq from all four groups of CAR- T cells, either without stimulation (baseline) or with 2 rounds of NALM6GL stimulation. For CAR-T cells that were cocultured with 2 rounds of NALM6GL cells, we sorted the pure CAR-T populations (CAR^+^; GFP^-^) before subjecting them to RNA sequencing (**Fig. S5a**). Principal component analysis (PCA) showed distinct group separation between control CAR, CAR-1CCT, CAR-2CCT and CAR-3CCT groups (**Dataset S1; Fig. S5b and S5c**). At baseline without NALM6GL stimulation, CAR-1CCT, CAR-2CCT and CAR-3CCT cells displayed reduced tonic signaling, as indicated by lower expression of inhibitory receptors including *LAG3, CD101, HHLA2, CD160, KLRB1* and *CD244* (**Fig. S5d**). High expression of these genes without antigen stimulation is consistent with antigen-independent activation of CARs, a hallmark of tonic signaling that can result in exhaustion^39, 40^. We further confirmed these findings at the protein level by flow cytometry analysis of LAG3, HHLA2 and CD101 (**Fig. S5e-g**).

Interestingly, from RNA-seq differential expression analysis of CAR-T cells that underwent repeated cancer stimulations, we observed that a single synthetic CCT substantially impacted the transcriptome. Furthermore, with increasing numbers of fused CCTs, more DE genes were identified. More specifically, compared CAR-1CCT with the control CAR-T cells, a total of 18 differentially expressed genes (DEGs) were found to be significantly upregulated and 15 genes were significantly downregulated (|log2FC| > 1; adjusted q < 0.01). CAR-2CCT cells showed 201 upregulated and 226 downregulated genes, and CAR-3CCT cells showed 516 upregulated and 483 downregulated genes (**Dataset S1; Fig. S5h**). Pathway analysis revealed a number of enriched differentially altered pathways (**Dataset S1; Fig. S5i**). For example, CAR-2CCT cells upregulated genes that are enriched in cell chemotaxis, immune cell / leukocyte migration and regulation of cytosolic calcium ion concentration, which are indicative of enhanced responsiveness of CAR-T cells to repeated tumor antigen stimulations (**Fig. S5i**). Among the commonly upregulated genes, *GZMK, CD244, CCR2, CCR5, CXCR6 and KLF2* are related to effector T functions, which were significantly upregulated in all three groups of CAR-T cells with CCTs (**Fig. S5j**). Notably, we also observed 20 upregulated DEGs unique to the CAR-2CCT cells, including *IL1R1, EPAS1, CCL2, RIPK2, KLRD1* and *IL24* (**Fig. S5j**). Among these, *IL1R1*^41^ and *EPAS1*^42^(encodes hypoxia-inducible factor 2a, HIF2a) were reported to directly enhance the effector function of adoptively transferred T cells against tumors. Together, these results demonstrate that CCT fusion can reduce tonic signaling of CARs at baseline while preserving high responsiveness of CAR-T cells to repeated antigen stimulations.

### Engineered CAR-T cells with monomeric or duplex CCT fusion exhibit superior *in vivo* therapeutic efficacy

To assess the *in vivo* performance of the synthetic CCT fusion CAR-T cells, we established a relapsed leukemia mouse model (**Fig. 6a**). In this model, NSG mice first underwent disease induction with 0.5 million of luciferase-expressing NALM6GL cells intravenously (i.v.) injected on day 0, and treated with 1 million CAR-T cells (i.v.) on day 4. To support *in vivo* CAR-T cell survival and better model physiologic T cell survival signals in NSG mice^43^, all mice were subcutaneously (s.c.) administered with 2.5µg human recombinant IL2 everyday starting on day 4 for 24 days. Using bioluminescence imaging of disease progression by the *In Vivo* Imaging System (IVIS), we found that all mice receiving CAR-T treatment had no detectable disease at day 7. We therefore modeled disease relapse by re-challenging with 0.5 million NALM6GL cells (i.v.) on day 12, when no luciferase signal was detected across all groups treated with CAR-T cells.

**Figure 6.**
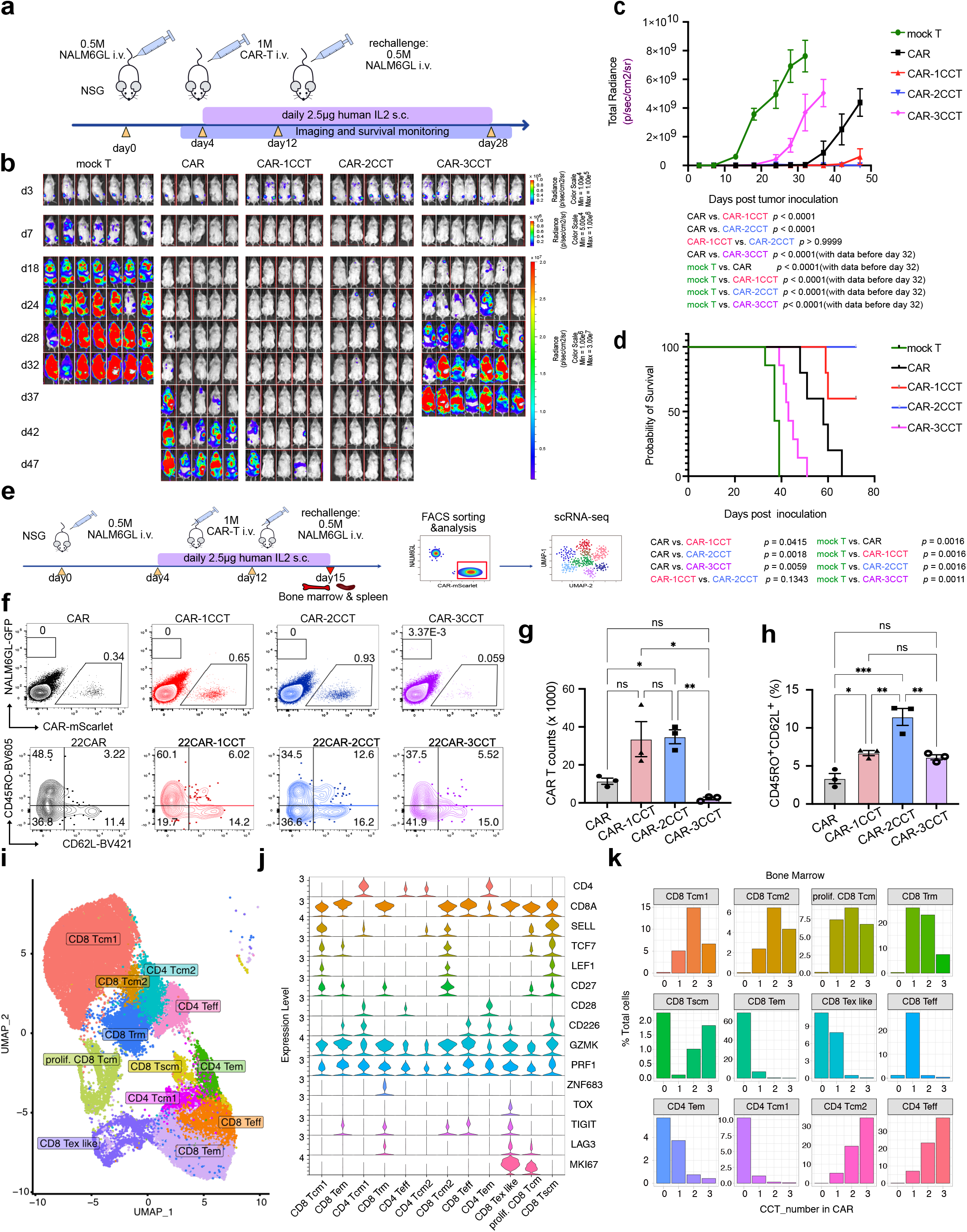
CAR-T with 1CCT or 2CCT fusion demonstrated improved anti-tumor efficacy in a relapsed leukemia mouse model with increased persistence and Tcm differentiation. **a.** Schematics of the *in vivo* experimental workflow used in **c-d**. NSG mice (6-10 weeks old) were inoculated with 0.5 million NALM6GL cells intravenously (i.v.) on day 0 and treated with 1 million CAR T cells (i.v.) on day 4. All mice were re-challenged with 0.5 million NALM6GL cells (i.v.) on day 12. 2.5µg human recombinant IL2 were administrated subcutaneously (s.c.) daily starting on day 4 for 24 days. Disease progression was monitored by bioluminescence imaging and survival analysis. **b.** Representative bioluminescence imaging of relapsed leukemia mouse model after CAR-T treatment. **c.** Quantification of **b** showing the tumor burden, as indicated by total radiance(p/sec/cm^2^/sr). Data are shown as mean ± s.e.m. Two-way ANOVA with Tukey’s multiple-comparisons test is used to assess significance. All comparisons with mock T group are done with data collected before day 32 (including day 32), when radiance of mice in mock T group is available. *p* values for comparisons are shown below the figure. **d.** Survival curve of NSG mice in (a). Mantel-Cox test is used to assess significance. *p* values comparisons are shown below the figure. **e.** Schematics of the workflow demonstrating the *in vivo* phenotyping of CAR-T cells. NSG mice (6-10 weeks old) were inoculated with 0.5 million NALM6GL cells intravenously (i.v.) on day 0 and treated with 1 million CAR-T cells (i.v.) on day 4. All mice were re-challenged with 0.5 million NALM6GL cells (i.v.) on day 12. 2.5µg human recombinant IL2 were administrated subcutaneously (s.c.) daily from day 4 to day 14. On day 15, bone marrow and spleen of each mouse were collected and processed for flow cytometry. Live mScarlet^+^ CAR-T cells were sorted out for scRNA-seq analysis. **f.** Representative flow cytometry results showing abundance of CAR-T cells and NALM6GL cells in the spleen (upper panel), and the CD45RO^+^CD62L^+^ central memory CAR-T cells in the bone marrow (lower panel) .**g-h**. Quantification of **f** showing CAR-T counts (**g**) and the percentage of CD45RO^+^CD62L^+^ central memory CAR-T cells (**h**). **i.** UMAP visualization of 39,151 single CAR-T cells based on their transcriptomes profiled by scRNA-seq. A total of 12 distinct clusters were identified and annotated. Tcm, T central memory cells; Teff, T effector cells; Tem, T effector memory cells; Trm, T residential cells; Tex like, T exhausted like cells; Tscm, T stem memory cells. **j.** Violin plots showing the expression levels of representative marker genes across 12 clusters. **k.** Relative proportions of each cell type in bone marrow identified by scRNA-seq, across four CAR-T groups. For all bar plots, data are shown as mean ± s.e.m. One-way ANOVA with Dunnett’s multiple-comparisons test is used to assess significance for figure (g) and (h). ns = *p*> 0.05, * = *p* < 0.05, ** = *p* < 0.01, *** = *p* < 0.001, **** = *p* < 0.0001.

By IVIS imaging, relapse was detected in mice treated with the control CAR-T as early as day 37, at which point no obvious luciferase signal was detected in mice from CAR-1CCT and CAR- 2CCT groups (**Fig. 6b and 6c**). CAR-1CCT cells efficiently controlled leukemia development in NSG mice, leading to significantly reduced tumor burden and significantly better survival than the control CAR-T cells (compared to control CAR, *p* < 0.0001 for tumor burden, *p* = 0.04 for survival) (**Fig. 6c and 6d**). CAR-2CCT cells had remarkably enhanced efficacy, in which all mice were in full remission against two cancer challenges, with 100% survival and nearly no detectable tumor burden (compared to the control CAR, *p* < 0.0001 for tumor burden, *p* = 0.0018 for survival) (**Fig. 6b-d**). Surprisingly, mice treated with CAR-3CCT cells had earlier relapses than those with CAR-T cells, indicating that triplex CCTs did not improve *in vivo* CAR-T efficacy over the control CAR- T cells. These data suggest that engineering CAR-T with monomeric or duplex CCTs significantly enhanced *in vivo* therapeutic efficacy against relapsed leukemia.

### CAR-2CCT cells strike the balance of persistence and central memory formation

Despite a higher relative survival rate observed in CAR-3CCT cells shortly after their adoptive transfer (**Fig. 5a-c**), CAR-3CCT cells fail to maintain long-term anti-tumor efficacy, suggesting that factors other than initial survival contribute to their weakened anti-tumor effects *in vivo*. To resolve this puzzle and gain a deeper understanding of the improved *in vivo* efficacy observed in CAR-T cells with 1CCT or 2CCT fusion, we performed immunological characterization of CAR- T cells in an *in vivo* model similar to that used in the efficacy measurement (**Fig. 6e**). In comparison to the control CAR-T cells, both CAR-1CCT and CAR-2CCT cells showed an upward trend in abundance in the spleen, with CAR-2CCT cells exhibiting a significant increase, whereas CAR- 3CCT cells were nearly undetectable. (**Fig. 6f and 6g**). Characterization of T cell memory phenotype showed that CAR-1CCT and CAR-2CCT cells had a notably higher fraction of central memory T cells (Tcm, CD45RO^+^;CD62L^+^), with CAR-2CCT cells having the highest Tcm% level; in contrast, CAR-3CCT cells did not have a significantly higher Tcm% compared to control (**Fig. 6f and 6h**). These data showed that CAR-T engineering with 1CCT and particularly 2CCT, but not 3CCT, enhanced *in vivo* persistence and central memory formation, in accordance with their superior anti-leukemia efficacy *in vivo*.

To systematically investigate the effect of CCT engineering on CAR-T cell phenotypes *in vivo*, we used single-cell RNA sequencing (scRNA-seq) to characterize the full spectrum of engineered CAR-T cells and their transcriptomic profiles. We isolated CAR-T cells from both spleen and bone marrow of the mice from the same leukemia mouse model (**Fig. 6e**). To minimize sampling bias, we pooled samples from 3 individual mice within the same group for all four groups (CAR, CAR- 1CCT, CAR-2CCT and CAR-3CCT). CAR-T cells were isolated from these pooled samples via fluorescence-activated cell sorting (FACS) based on the expression of mScarlet (**Fig. S6a**). We mapped the transcriptomes of 39,151 single CAR-T cells via Uniform Manifold Approximation and Projection (UMAP), and identified 12 distinct clusters (**Fig. 6i and Dataset 2**). We annotated those clusters based on their expression profile of known immune-cell markers (**Dataset 2**). When annotating based on *CD4* and *CD8A* expression, we observed that CD8 CAR-T and CD4 CAR-T cells comprised 8 and 4 clusters, respectively (**Fig. 6i**). Heterogeneity within these populations was further resolved based on the expression of naïve/memory markers (*CD62L, TCF7, LEF1, CD27 and CD28*), effector/cytotoxicity associated markers (*GZMK, PRF1*), activation/exhaustion markers (*CD226, TOX, TIGIT and LAG3*), tissue residency marker (*ZNF683*) and proliferation marker (*MKI67*) (**Fig.6j, Fig. S6b and Dataset 2**). In terms of tissue distribution, samples isolated from bone marrow comprised a higher level of heterogeneity than those from spleen, as more diverse T cell states were identified in CAR-T populations from bone marrow than spleen (**Fig. S6c**).

scRNA-seq of the CCT engineered CAR-T cells (CAR-1CCT, CAR-2CCT and CAR-3CCT) revealed significant changes in cell subset composition compared to the CAR control group in the bone marrow (**Fig. S6d**), but not spleen (**Fig. S6e**). We observed an increase of CD8 Tcm1, CD8 Tcm2, proliferating (prolif.) CD8 Tcm and CD8 tissue resident T (CD8 Trm) cells in CAR-1CCT and CAR-2CCT particularly, relative to the control CAR group (**Fig. 6k**). Consistent with the flow cytometry results, we observed major differences in the abundance of Tcm cells between CCT engineered CAR-T cells and control CAR-T cells (**Fig. 6i, Fig. S6d**). CAR-2CCT cells from bone marrow had the highest proportion of CD8 Tcm cells among all groups (**Fig. 6k and S6d**). We further analyzed gene DEGs between CAR-2CCT and CAR-3CCT, which had the most distinct *in vivo* efficacy. In most T cell clusters, the number of DEGs are small (**Supplemental Dataset 2**). We noticed that in the bone marrow CD4 Teff cluster, effector makers like *KLRB1, KLRK1, KLRG1, NKG7, GZMK* and *CTSW* were significantly upregulated in the CAR-2CCT compared with CAR-3CCT group (**Fig. S6f**). This suggests that CD4 Teff cells from the CAR-2CCT group have a more activated and cytotoxic phenotype compared to those from the CAR-3CCT group. Taken together, these data suggest that the improved *in vivo* efficacy achieved by CAR-1CCT and CAR-2CCT cells is linked to their heightened persistence and ability to differentiate into Tcm cells when faced with repeated challenges.

## Discussion

Cell-based therapeutics have demonstrated great promise with six CAR-T cell therapies currently approved by the FDA and over one thousand active clinical trials involving cell therapies (clinicaltrials.gov). Despite these advances, durable remissions are only observed in a limited number of patients^44^. Improving the clinical efficacy of CAR-T cell therapy thus necessitates the development of new strategies to prevent relapse. By harnessing the unique dynamic features of the CTLA-4 cytoplasmic tail (CCT), we developed CAR-T cells in which the CAR is fused to varying numbers of CCTs. Mechanistically, CCT fusion altered the dynamics of CAR molecules by accelerating CAR endocytosis, degradation and recycling, processes that we observed were infrequently occurring with native CARs. This in turn led to reduced CAR-T tonic signaling and dampened T cell activation and inflammatory cytokine production. Additionally, CCT fusion further limited trogocytosis and cancer antigen loss, ameliorating the potential for subsequent CAR-T fratricide. In turn, CAR-1CCT and CAR-2CCT cells showed improved survival and persistence in the context of a leukemia mouse model, with enhanced anti-cancer functionality upon repeated cancer stimulation, increased *in vivo* persistence, and enrichment for Tcm differentiation (**Fig. S7**).

Trogocytosis is well-documented process that mediates surface molecule transfer from target cells to many immune populations such as T ^11^, B ^12^ and NK ^13^ cells. Recently, CAR mediated trogocytosis was reported to impair the anti-tumor efficacy of both CAR-NK ^45^ and CAR-T ^15, 16^ cell-based therapies. However, to date there are no established strategies that can be generally applied to reduce CAR-mediated trogocytosis. CCT fusion approach introduces a new molecular feature to the CAR, enabling dynamic control of surface CAR expression at the protein level and effectively reducing CAR-mediated trogocytosis. Importantly, by adjusting the number of CCT fusions, we demonstrated quantitative regulation of CAR-mediated trogocytosis (**Fig. 3**). Our study therefore builds on previous work showing that the frequency of trogocytosis is impacted by the density of target antigen^46^ and the affinity of CAR^16^. To our knowledge, our study is the first to show that the molecular dynamics of CARs and CAR-mediated trogocytosis can be quantitatively regulated by adjusting the number of CCT fusions. This approach involves engineering the distal cytoplasmic end of the CAR, which might be further tested and applied to receptors with different antigen specificities. We show here that the properties of CCT fusion are also applicable to CD19-specific CARs, suggesting that it is not limited to one type of CAR (CD22 CAR). In the future, CCT fusion might also be tested for the control of trogocytosis mediated by other cellular therapies such as CAR-NK cells, as NK cells are reported to have similar regulation machinery to CTLA-4 endocytosis in CD4^+^ T cells ^47, 48^.

Besides the regulation of trogocytosis, CCT fusion effectively improved CAR-T survival *in vitro* and *in vivo* (as shown above in **Fig.1d-e, Fig. 4b-d**), while also leading to reduced apoptosis (**Fig. 4h-i**). While the reduction in trogocytosis and potentially fratricide among CAR-T cells with CCT likely contributes to this phenomenon, other possible explanations such as increased proliferation (**Fig. 4e-f**) and reduced cell activation-induced death are also plausible explanations.

Interaction strength between CAR and its cognate antigen is a fundamental determinant of CAR- T function ^49^. In line with this, our data show that regulating CAR signaling strength with CCT fusion prevents CAR-T cells from excessive stimulation and enables durable response against tumor cells. CAR signaling needs to be tightly regulated to achieve the fine-tuned balance of sufficient signaling for effective tumor antigen detection and CAR-T action, while also circumventing the detrimental effects seen in CARs with an excessively high affinity or expression ^8, 50, 51^. This resonates in our results with CAR-3CCT cells, as these cells exhibited the best killing capability under repeated stimulation *in vitro* yet performed worse than the control CAR-T cells *in vivo*. One possible explanation is that the surface availability of CAR needed for optimal signaling strength is different *in vitro* vs *in vivo*. *In vitro* co-culture assays have artificially high density and close proximity of CAR-T cells to tumor cells; in this context, reduced CAR surface expression (as seen in CAR-3CCT) is potentially advantageous by sufficiently mediating tumor cell killing while mitigating drawbacks such as exhaustion, activation-induced cell death, and fratricide. However, CAR-CCT cells are more diluted in the *in vivo* setting, and perhaps higher CAR surface expression and signaling strength is crucial to overcome the lower density of CAR- CCT cells *in vivo*. CAR-2CCT cells may land in a local optimum for *in vivo* efficacy with balanced cellular features. More generally, our study highlights the critical importance of *in vivo* studies to titrate the signaling strength induced by CARs for sustained anti-tumor responses, as *in vitro* co-culture assays often can not recapitulate such complexities.

Other studies have proposed approaches to modulate CAR signaling strength, such as transient cessation of CAR signaling with a multi-kinase inhibitor ^7^, lower affinity CARs ^8^, and regulated promoters ^52, 53^. Distinct from these studies, CCT fusion, with the merit of simplicity, enables tunable regulation of CAR surface availability without additional chemicals, without the need to engineer new scFVs of varying affinity. CCT fusion effectively regulates CAR availability at the protein level, in certain cases advantageous over transcriptome level regulation (e.g. inducible promoters) because it provides direct control over the amount of CAR protein present, as the relation between transcriptional levels and downstream protein levels is often complex. CCT fusion also opens up a potential avenue to generate CARs with self-regulation according to antigen stimulation level. For instance, fused CCT(s) could be cleaved in a regulatable manner by introducing a protease module that is responsive to antigen stimulation, thus enabling control over the timing and duration of CAR expression. This self-regulation would allow for flexible and dynamic self-titration of CAR signaling, which we envision will be valuable in optimizing efficacy while also limiting toxicity.

In conclusion, our study demonstrates that fine-tuning CAR dynamics with CCT fusion can improve the anti-tumor efficacy of CAR-T-cell therapy, illuminating an orthogonal strategy for the development of more effective CAR-T-cell therapies. Future studies can explore the generalizability of this tail fusion strategy, including different protein tails, different cancer types, different CARs and different types of therapeutic immune cells.

## Institutional approval

All recombinant DNA work was performed under the guidelines of Yale RBC and EHS with approved protocols (Chen-15-45 and Chen-20-18). All animal work was performed under the guidelines of Yale University Institutional Animal Care and Use Committee (IACUC) with approved protocols (Chen-2015-20068; Chen-2018-20068; Chen-2021-20068).

## Discussion and Support

We thank Drs. Krause, Isufi, and Bersenev for discussions on cell therapy. We thank all members in Chen laboratory, as well as various colleagues and core facility staff at Yale for assistance and/or discussion. We thank the Yale West Campus Imaging Core for the support and assistance in this work. We thank Dr. R Glen for helpful proofreading on the manuscript.

## Methods

### Vector construction

All lentivirus plasmids were generated by inserting target coding sequences into lentivirus transfer plasmid backbone (Addgene, #75112). Specifically, the pLenti-EFS-Flag-scFV(m971)-CD28- 41BB-CD3zeta-T2A-mScarlet (CD22-CAR) was generated by amplifying the sequence from a gblock synthesized by IDT, and inserting it into the lentiviral backbone using Gibson Assembly. Afterwards, to generate CAR-CCT constructs, 1CCT, 2CCT and 3CCT were inserted right before the T2A sequences, by amplifying 1-3 CCTs from the synthesized sequences with three tandem CCTs. CD19-CAR and CD19-CAR-CCT constructs were generated by replacing the m971 scFv with the FMC63 scFv from above CD22-CAR constructs. Similar methods were used to generate pLenti-EFS-CD22-GGGGS-mTagBFP and pLenti-EFS-CD22-GGGGS-eGFP constructs.

### Cell culture

293T, NALM6 and Jurkat cells were purchased from ATCC. Human PBMCs were purchased from Stemcell. NALM6 expressing GFP-luciferase (NALM6GL) was previously generated in lab^54^. 293T cells were cultured in DMEM (Gibco) media supplemented with 10 % FBS (CORNING) and 200 U/mL penicillin-streptomycin (Gibco), hereafter referred to as cDMEM. NALM6 cells and Jurkat cells were cultured in RPMI-1640 (Gibco) media supplemented with 10% FBS and 200 U/mL penicillin-streptomycin, hereafter referred to as cRPMI. Human PBMCs were cultured in X-VIVO^TM^ 15 media (Lonza) supplied with 5 % human AB serum (MP Biomedical) and 10ng/mL human IL-2 (Peprotech), hereafter referred to as cX-VIVO. All cells were grown at 37°C and 5% CO_2_ with saturating humidity.

### Lentivirus production

One day before transfection, 40 million 293T cells (passage number less than 20) were seeded into a 150mm dish. The next day, old medium was replaced with pre-warmed fresh cDMEM. For each dish, 20ug transfer plasmid, 10ug packaging plasmid psPAX2 (Addgene) and 5ug envelop plasmid pMD2.G (Addgene) were diluted with 1mL plain DMEM. In a separate tube, 87.5µl LipoD293 was diluted with 1mL plain DMEM and then added to the diluted DNA mixture. This transfection mix was then briefly vortexed and incubated at room temperature for 15 minutes before being added to the 293T cells. Lentivirus was harvested 48 hours post transfection by collecting the supernatant of transfected cells. The supernatant was then spun down at 3000g for 15 minutes to get rid of cell debris. Lentivirus was concentrated by adding 40% (w/v) PEG8000 directly to the supernatant to final concentration of 8% (w/v) PEG8000, and then incubated at 4°C overnight. The next day, the lentivirus was spun down at 1500g for 30 minutes, pegylated viral pellet was resuspended with 1mL fresh cRPMI or cX-VIVO medium. Virus was then stored at -80°C before usage.

### Generation of stable cell lines

All stable cell lines used in this study were generated by lentiviral infection. Briefly Jurkat, NALM6 or NALM6GL cells were spin-infected with lentiviruses carrying corresponding constructs at 2000rpm (∼900g), 37 °C for 2 hours. Infected cells were then purified based on reporter gene (GFP, BFP or mScarlet) expression by flow cytometer. NALM6-CD22-GFP cells were generated through transduction with a lentiviral vector that expressed full-length human CD22 and eGFP linked by a GGGGS linker. NALM6GL-CD22-BFP cells were generated using a similar lentiviral construct, where eGFP was swapped for mTagBFP in NALM6GL cells. CAR- Jurkat cells were generated by transducing Jurkat cells using the same lentiviral constructs showed in **Fig. 1a**.

### Human T cell isolation and culture

Human CD3 T cells were isolated directly from PBMC samples by using magnetic positive selection. Briefly, 15 million PBMCs were labeled with anti-human CD3-biotin (Biolegend, 1:100) in MACS buffer (PBS supplemented with 0.5% BSA and 2mM EDTA) at 4°C for 10 minutes. After being washed with MACS buffer, PBMCs were resuspended with 120µL MACS buffer and then stained with 30uL of anti-biotin beads (Miltenyi) at 4°C for 15 minutes. After a final wash with MACS buffer, labeled CD3 T cells were separated by using MACS LS Columns (Miltenyi) and MACS Separators (Miltenyi) according to the manufacturer’s instructions. Purified CD3 T cells were cultured in cX-VIVO medium and stimulated with Dynabeads® Human T-Activator CD3/CD28 (Thermo) at a bead-to-cell ratio of 1:1 for 24 hours before being infected with lentiviruses.

### Human T cell transduction with lentivirus

Human CD3 T cells activated for 24 hours were collected, and resuspended with concentrated lentiviruses, supplemented with 8µg/mL polybrene, at a concentration of 1 million/mL. Cells were plated in 24-well plate and spun at 2000rpm (∼900g) for 2 hours at 37°C. Immediately after the spin, the virus supernatant was aspirated and replaced with fresh cX-VIVO medium. Cells were split 1 to 2 every other day and transgene expression was checked 4 days after infection by flow cytometry. CAR-T cells were sorted on day 7 based on mScarlet expression and expanded in vitro for another 7 days before being used for *in vitro* and *in vivo* experiments.

### Antibody and staining for flow cytometry

All antibodies used in this study are listed in the Antibodies section of the Nature Portfolio Reporting Summary. Cell surface antigens were stained with indicated antibody cocktails in MACS buffer on ice for 15 minutes as indicated in the figure legend. To stain endocytic CAR or cycling CTLA-4, CAR-T cells were stained using anti-flag-BV421(Rat-IgG2a, Biolegend) or anti-CTLA-4-APC (Mouse IgG2a, Biolegend) at 37°C for 1hour, followed by washing with ice old MACS buffer. Those cells were then stained with anti-Rat-IgG2a-Alexaflour647 (Biolegend) or anti-mouse IgG2a-BV421(Biolegend) at 4°C for 15 minutes before running on the flow cytometer. To quantify apoptosis, cells were stained with Annexin V Pacific-blue conjugates (ThermoFisher) in Annexin V binding buffer (BD) at room temperature for 15 minutes, cells were then immediately analyzed by flow cytometer. LIVE/DEAD Fixable Near-IR Dead Cell Stain (Invitrogen) was included for all staining to exclude unspecific staining of dead cells. Samples were collected or sorted on a 4-laser (405nm, 488nm, 561nm and 640nm) Aria II cell sorter (BD). For sorting experiments, sorting purity was checked after every sort to make certain it was higher than 95%. All flow cytometry data were analyzed using FlowJo software 10.8.0 (BD).

### *In vitro* CAR T killing assay

Variable amounts of purified CAR T cells were cultured with fixed amount of NALM6GL cells at an E/T ratio of 1/2, 1/4 or 1/8 in a 96-well U bottom plate. 24 hours later, cells were either collected for flow cytometry analysis or re-stimulated with the same amount of NALM6GL cells as used in the first round coculture. 24 or 48 hours later (indicated in the figure legend), NALM6GL cells and CAR-T cells were stained with indicated antibody cocktails. Cell counting beads (Biolegend) were added to all samples for absolute cell number measurement. For multiplex cytokine detection, the supernatant from cocultures was collected and measured by LEGENDplex™ Human CD8/NK Panel (Biolegend) according to the manufacturer’s instructions.

### Proliferation assay

Purified CAR-T cells were stained in PBS with 10µM Cell Proliferation Dye eFluor™ 450 (ThermoFisher) at 37°C for 5 minutes. Cells were then washed three times with cRPMI medium and resuspended in cX-VIVO medium at a final concentration of 0.5 million/mL. 200µL cells were seeded into a 96-well U bottom plate. 4 days later, eFluor450 dilution was measured by flow cytometry.

### Measurement of phosphorylation and cytokine production

To measure phospho-ERK, CAR-T cells were first starved overnight in serum free X-VIVO medium, and stained for LIVE/DEAD Fixable Near-IR Dead Cell Stain (Invitrogen). After washed with ice old PBS, those CAR-T cells were incubated with 2µg/mL biotinylated human CD22 (ECD) on ice for 30 minutes. Signal transduction was induced by crosslinking with 10µg/mL streptavidin (Biolegend) at 37°C for 5minutes. Crosslinking was stopped immediately with the Phosflow™ Fix Buffer I (BD) and fixed at 37°C for 10 minutes. After this, cells were permeabilized with BD Phosflow™ Perm Buffer III on ice for 30 minutes, followed by Anti- ERK1/2 (pT202/pY204)-Alexaflour647 (Biolegend) staining at room temperature for 30 minutes. For the intracellular IFNg and TNFa staining, CAR-T cells were cocultured with NALM6GL cells at indicate E/T ratio for 4 hours with the presence of 1x Brefeldin A (Biolegend). Cells were first stained with the Live/Dead-NearIR dye before fixed, permeabilized and stained with BD Cytofix/Cytoperm™ Buffer System according to manufacturer’s suggestion.

### Measurement of recycling CAR

The method used for quantification of recycling receptors over time has been described by others^32^. Briefly, CAR-T cells were first stained with anti-Flag (Biolegend) at 37°C for 1 hour, and washed with ice cold MACS buffer. The anti-Rat IgG2a-Alexaflour647 secondary antibody either stained at 4°C for 15 minutes (baseline) or at 37°C for 30 minutes or 60 minutes in order to see an increase in the secondary antibody signal when compared to the baseline. The increase in the percentage of cells stained positive for the secondary antibody was quantified by flow cytometry as an indication of recycling.

### Measurement of recycling CAR or CTLA-4

The method used for quantification of recycling receptors over time has been described by others^32^. Briefly, CAR-T cells were first stained with anti-Flag antibody (Biolegend) or anti-CTLA-4 antibody (Biolegend) at 37°C for 1 hour, and washed with ice cold MACS buffer. The secondary antibody either stained at 4°C for 15 minutes (baseline) or at 37°C for 30 minutes or 60 minutes in order to see an increase in the secondary antibody signal when compared to the baseline. The increase (fold change) in the percentage of cells stained positive for the secondary antibody was quantified by flow cytometry as an indication of recycling.

### Cycloheximide chase assay

The stability of CAR was quantified by treating CAR-T cells with 50µg/mL cycloheximide (Sigma) at 37°C for up to 4 hours. Total CAR expression was quantified by staining intracellular flag expression using BD Cytofix/Cytoperm™ Buffer System according to manufacturer’s suggestion.

### CAR degradation with antibody feeding

The method used for quantification of receptor degradation over time has been described by others^32^. Briefly, CAR-T cells were incubated with anti-Flag antibody (Biolegend) at 37°C for 1 hour. Cells were then washed and incubated at 37°C for up to 4 hours with or without 10µM bortezomib (Sigma) or 10nM BafA1. Cells were then fixed and permeabilized with BD Cytofix/Cytoperm™ Buffer System, and stained with Anti-Rat-IgG2a-Alexaflour647 before analyzed by flow cytometry.

### Quantification of CD22 transfer by confocal microscopy

About 10,000 Purified CAR-T cells that had performed trogocytosis (mScarlet^+^ BFP^+^) were seeded onto µ-Slide 8-Well glass bottom chamber (Ibidi) that had been previously treated with poly-L-lysine. Cells were then stained with deep red CellMask^TM^ Plasma Membrane dye (ThermoFisher) at 37°C for 5 minutes according to the manufacturer’s instruction. Cell were imaged by Leica Stellaris 8 FALCON with the 40× objective (oil). Representative images were exported using ImageJ (https://imagej.nih.gov/). The percentage of CD22 membrane colocalization was analyzed by a customized MATLAB (Mathworks, Natick, MA) script. Specifically, Images of BFP tagged CD22 antigen and CellMask stained membrane were first binarized with a global threshold for pixel intensity (10 for antigen and 15 for membrane) to classify each pixel into either foreground pixel or background pixel. Cells appear as a ring in membrane mask and all closed rings were filled to generate a binarized image where foreground pixels represent intact cells (cell mask). Cytosol mask was obtained by subtracting membrane mask from cell mask. Antigen mask then underwent a size filtering to remove objects with too few (<3 pixels) or too many pixels (>300 pixels) which mainly originates from impulse noise and dust. Cell mask underwent a similar size filtering (<100 pixels and >1000 pixels) to remove objects originating from impulse noise, broken cells and clusters of cells such that the filtered cell mask only contains well isolated single cells. Each single cell in the filtered cell mask was then discarded/selected for final colocalization calculation based on two quality control indexes: 1) cytosol percentage; 2) number of contained antigen pixels. Too low cytosol percentage (<40%) might be caused by endocytosis of the membrane staining dye into cells and can lead to unreliable membrane segmentation. Therefore, cells with too low cytosol percentage were excluded from the analysis. Too few antigen pixels (<20 pixels) lead to significant uncertainty in estimation of colocalization percentage and thus were also excluded from further analysis. Finally, for all the remaining cells, antigen-membrane colocalization percentage was calculated by the following formula where 𝐼_𝑎𝑛𝑡𝑖𝑔𝑒𝑛-𝑐𝑦𝑡𝑜𝑠𝑜𝑙_, 𝐼_𝑎𝑛𝑡𝑖𝑔𝑒𝑛-𝑐𝑒𝑙𝑙,_ stands for intensity of each pixel classified as foreground in both antigen and cytosol mask, respectively. 𝐼_𝑎𝑛𝑡𝑖𝑔𝑒𝑛 𝑏𝑎𝑐𝑘𝑔𝑟𝑜𝑢𝑢𝑛𝑑_ stands for mean intensity of background pixels in antigen image (2.5 used for all analysis).

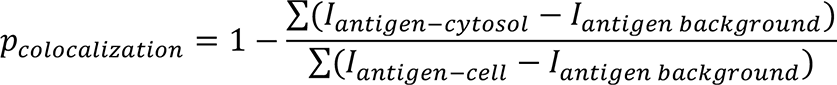

### Time-lapse live cell imaging of trogocytosis and fratricide

40, 000 CAR-T cells and NALM6GL-CD22-BFP cells were seeded at an E to T ratio of 2:1 into a µ-Slide 8-Well glass bottom chamber (Ibidi) that had been previously treated with 7.5µg/cm^2^ poly-L-lysine (Sigma). 5nM SYTOX^TM^ Deep Red Nucleic Acid Stain (ThermoFisher) was supplemented into the culture to allow for dead cell staining. Cells were imaged with Leica Stellaris 8 FALCON with the 40× objective (oil) in the live cell imaging chamber at 37°C and 5% CO2. A 4 x 4 field-of- view was taken at a scan interval of every 3 to 5 minutes for up to 8 hours. Images were processed and exported through ImageJ (https://imagej.nih.gov/).

### Trogocytosis assay

CAR-T or CAR-Jurkat cells were coculture with NALM6GL, NALM6GL-CD22-BFP or NALM6-CD22-GFP cells for indicated time at a fixed E to T ratio, as specified in the figure legend. Trogocytosis of CD22 by CAR-T cells were quantified by measuring CD22 using antibody, BFP or GFP signal. For trogocytosis inhibition, as indicated in the figure legend, CAR T cells were pre-treated with 1 μM latrunculin A (Sigma-Aldrich) at 37 °C for 15 min before co-incubation with target cells.

### Degranulation assay

CAR-Jurkat cells were cocultured with NALM6-CD22-GFP at an E/T ratio of 1:1 for 4 hours to allow for CD22-GFP transfer from NALM6 to CAR-Jurkat cells. Both Trog^+^ CAR-Jurakt and Trog^-^ CAR-Jurkat cells were sorted out based on their expression of CD22-GFP using an AriaII sorter (BD). Those cells were then used as target cells and cocultured for 2 hours with fresh CAR- T cells that had been labeled with 10nM eFluor450 to allow for differentiation between CAR-T and CAR-Jurkat cells. Variable CAR-T to CAR-Jurkat ratios were used as indicated in the figure legend. Anti-CD107-APC (Biolegend) and 1x Monensin (Biolegend) were added into the culture during the incubation. Cells were then stained with LIVE/DEAD Fixable Near-IR Dead Cell Stain (Invitrogen) before analyzed by flow cytometry.

### Bulk RNA sequencing and data processing

To profile baseline differences, purified CAR-T cells expanded in vitro for 14 days were collected. To profile differences under repeated stimulations, CAR-T cells expanded in vitro for 14 days were cocultured with NALM6GL cells at E/T = 1/2, followed by a second round of NALM6GL cells stimulation 24 hours after the initial coculture. mScarlet^+^ CAR-T cells were then sorted 72hrs after the initial coculture. These purified CAR-T cells were lysed for bulk RNA extraction using RNeasy Plus mini isolation kit (Qiagen). Library preparations were performed using a NEBNext® Ultra™ RNA Library Prep Kit for Illumina according to manufacturer’s instruction. Samples were barcoded and multiplexed using NEBNext® Multiplex Oligos for Illumina® (Index Primers Set 1). Libraries were then sequenced with Novaseq (Illumina). Fastq files generated by sequencing were aligned and mapped using STAR^55^ with the human genome assembly version GRCh38. Differential expression was analyzed using DEseq2^56^. *p* (adj) < 0.01 and |log2FoldChange| > 1 were set as the cutoff for determining differentially expressed genes, which were selected for over representation analysis (ORA) using R package enrichplot^57^. Visualizations of differentially expressed genes, such as volcano plots and heatmaps, were generated using standard R packages such as pheatmap and VennDiagram.

### Measurement of relative CAR-T survival

CAR, CAR-1CCT, CAR-2CCT and CAR-3CCT cells stained with 1µM eFluor450 were mixed with CAR-T cells stained with 10µM eFluor450 at 1:1 ratio. For *in vitro* measurement, these mixed cells were then cultured with or without NALM6GL stimulation at an E/T ratio of 1:2 for 24 hours. For *in vivo* measurement, NSG mice (female, 6-10 weeks old) were first inoculated with 1 million NALM6GL cells intravenously (i.v.) on day 0. Two million of these labeled cell mixture was transferred to the leukemia NSG models on day 4. One day post CAR-T transfer, bone marrow samples were collected and processed into single cell suspension according to methods mentioned above. Cells were stained using LIVE/DEAD Fixable Near-IR Dead Cell Stain (Invitrogen) and anti-human CD22 APC (Biolegend) before being analyzed by flow cytometer. Flow cytometry was used to determine the relative percentage of eFluor450^low^ and eFluor450^high^ population. Relative survival of CAR T cells. And it was calculated by following equation: Relative survival (%) = [(l_w_ : h_w_)-(l_wo_ : h_wo_)]/ (l_wo_ : h_wo_)*100% , in which l_w_ and h_w_ stands for percentage of eFluor450 low and high population, respectively, with NALM6GL cells stimulation (*in vitro* assays) or after transfer (*in vivo* assays), while l_wo_ and h_wo_ stands for percentage of eFluor450 low and high population, respectively, without NALM6GL cells stimulation (*in vitro* assays) or before transfer (*in vivo* assays).

### *In vivo* evaluation of CAR-T performance

NOD.Cg-Prkdc^scid^ Il2rg^tm1Wjl^/SzJ (NSG) mice were purchased from the Jackson Laboratory and bred in house. NSG mice (female, 6-10 weeks old) were inoculated with 0.5 million NALM6GL cells intravenously (i.v.) on day 0 and treated with 1 million CAR-T cells (i.v.) on day 4. Relapse was modeled by re-challenging all mice with 0.5 million NALM6GL cells(i.v.) on day 12. To better model the clinical response to CAR-T cells in NSG mice^43^, 2.5µg human recombinant IL2 (Peprotech) was administered subcutaneously (s.c.) everyday starting on day 4 for 24 days. Disease progression was monitored by bioluminescence imaging and survival analysis. For the in vivo phenotyping of CAR-T cells, 2.5µg human recombinant IL2 was administered s.c. every day from day 4 to day 14. All mice were sacrificed on day 15. Spleen and bone marrow tissues were collected and processed into single cell suspension. For preparations, spleens were prepared by mashing them through 100um filters. For bone marrow isolation, femur with both ends cut by scissors was isolated, and bone marrow was flushed out with 2mL cRPMI using a 25G needle. For both spleen and bone marrow suspensions, red blood cells were lysed with ACK Lysis Buffer (Lonza), incubated for 2 minutes at room temperature, and washed with cRPMI. Lymphocytes were poured through a 40µm filter before staining with antibody cocktail. Cell counting beads (Biolegend) were added to all samples for absolute cell number measurement by Aria II cell sorter (BD).

### scRNA-seq library preparation and sequencing

Live CAR-T cells were sorted based on mScarlet expression from both the spleen and bone marrow samples processed as described above. For all four groups (CAR, CAR-1CCT, CAR-2CCT and CAR-3CCT), samples from 3 individual mice within the same group were pooled together to minimize sampling bias. Sorted cells were washed with PBS, and cell number and viability were assessed by trypan blue (Lonza) staining. About 2,000 to 10,000 purified mScarlet^+^ cells were used for scRNA-seq library preparation using Chromium Next GEM Single Cell 5’ Reagent Kits V2 (10x Genomics) according to manufacturer’s instructions. The single cell libraries were sequenced by NovaSeq 6000 (Illumina) with 2x150 read length.

### scRNA-seq data analysis

Analysis of scRNA-seq was performed using standard pipelines with custom codes. In brief, raw FASTQ data were pre-processed with Cell Ranger v6.0.1. The processed data matrices were then analyzed and visualized using the Seurat v4 package^58^. More specifically, the dataset was filtered to retain cells with < 10% mitochondrial counts and 200-2,500 unique expressed features^59^. The dataset was then log-normalized, scaled using the reciprocal-PCA dimensional reduction with 2000 anchors^60^. And dimensional reduction was performed by uniform manifold approximation and projection (UMAP)^61^ using the first 10 dimensions from PCA, which were chosen by the inflection point of an elbow plot. Cells were clustered in low-dimensional space by generating a shared nearest neighbor (SNN) graph (k = 20, first 10 PCs). Each cell cluster was annotated by specific type of T cells using canonical marker genes. An empirical resolution (0.2), followed by a second-step resolution for sub-clustering, was chosen for better separation of CD4 and CD8 T cell populations.

### Schematic illustrations

Schematic illustrations were created with Affinity Designer.

### Replication, randomization, blinding and reagent validations

Replication: Biological or technical replicate samples were randomized where appropriate. In animal experiments, mice were randomized by littermates and cages.

Sex as biological variable: Both male and female animals were used for experiments.

Blinding: *In vitro* CAR-T killing assay experiments were blinded by anonymized labeling. In such experiments, the group identity is unknown to data collector. Other experiments are not blinded. Resource validations: Commercial antibodies were validated by the vendors and re-validated in house as appropriate.

Authentication: Cell lines were authenticated by original ^8^vendors and re-validated in lab as appropriate by morphological examination and/or genetic verification.

Contamination check: All cell lines tested negative for mycoplasma. Misidentified cell lines: No commonly mis-identified cell lines involved.

### Code availability

The code used for data analysis and the generation of figures related to this study are available from the corresponding author upon reasonable request.

### Data and materials availability

All data generated or analyzed during this study are included in this article and its supplementary information files. Source data and statistics are provided in an excel file of **Source data and statistics**. Processed data for genomic sequencing and gene expression are provided as processed quantifications in **Supplementary Datasets**. Genomic sequencing raw data are being deposited to NIH Sequence Read Archive (SRA) and/or Gene Expression Omnibus (GEO), with pending accession numbers. Original cell lines are available at commercial sources listed in supplementary information files. Other relevant information, data or materials are available from the corresponding author upon reasonable request.

## Supplemental Figure Legends

**Figure S1.**
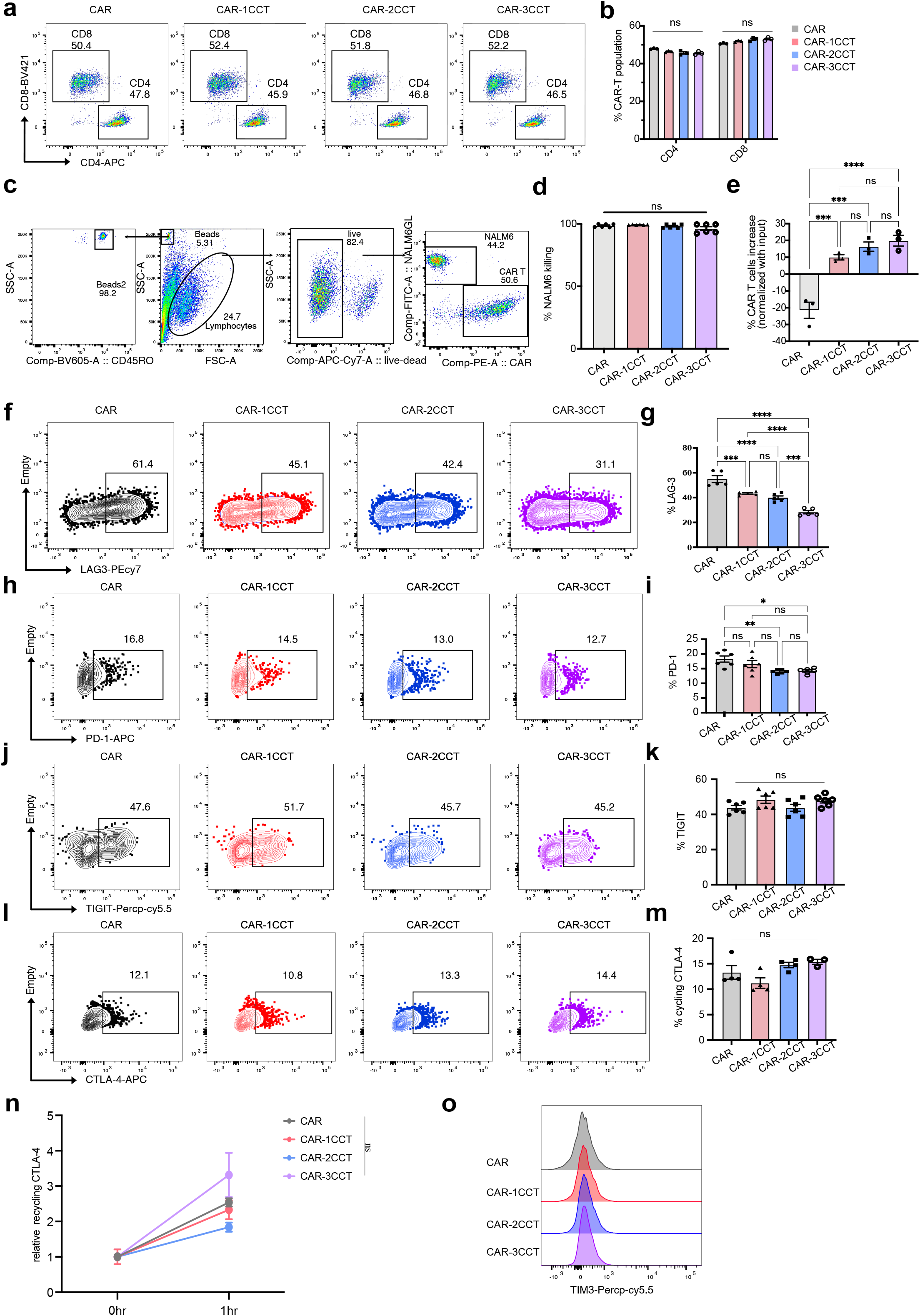
Characterization of CAR-T cells with CCT fusion after one-round stimulation with NALM6GL cells. **a.** Representative flow cytometry results showing CD4 and CD8 staining of CAR-T cells 7 days after transduced with CAR-T constructs. **b.** Quantification of **a.** showing the CD4 and CD8 percentage of CAR-T cells. **c.** Representative flow cytometry results showing the gating strategy used in Fig. 1 for *in vitro* CAR-T and NALM6GL cells coculture. **d.** Quantification of NALM6 killing by CAR-T cells. Purified CAR-T cells were cocultured with one round of NALM6GL cells at E/T = 1/2 for 24 hours. Cells were then collected and analyzed by flow cytometry. **e.** Quantification of the increases in CAR T cell counts after one round of NALM6GL stimulation. Purified CAR T cells were cocultured with one round of NALM6GL cells at E/T = 1/2 for 24 hours. Cells were then collected and analyzed by flow cytometry. CAR-T cells increase (%) is calculated by following equation: % CAR-T cells increase = (n_w_-n_wo_)/ now * 100%, in which n_w_ and n_wo_ stands for the number of CAR-T cells with or without (input) NALM6GL coculture, respectively. **f-o**. Representative flow cytometry analysis and quantification of LAG-3 (**f-g**), PD-1(**h-i**), TIGIT(**j-k**), cycling CTLA-4(**l-m**), recycling CTLA-4 (**n**) and TIM3 (**o**) on CAR-T cells after stimulated with NALM6GL cells at E/T = 1/2 for 24 hours. For all bar plots, data are shown as mean ± s.e.m. One-way ANOVA with Dunnett’s multiple-comparisons test is used to assess significance for figure (**d**), (**e**), (**g**), (**i**), (**k**), and (**m**). Two-way ANOVA with Tukey’s multiple-comparisons test is used to assess significance for figure (**b**). ns = *p*> 0.05, * = *p* < 0.05, ** = *p* < 0.01, *** = *p* < 0.001, **** = *p* < 0.0001.

**Figure S2.**
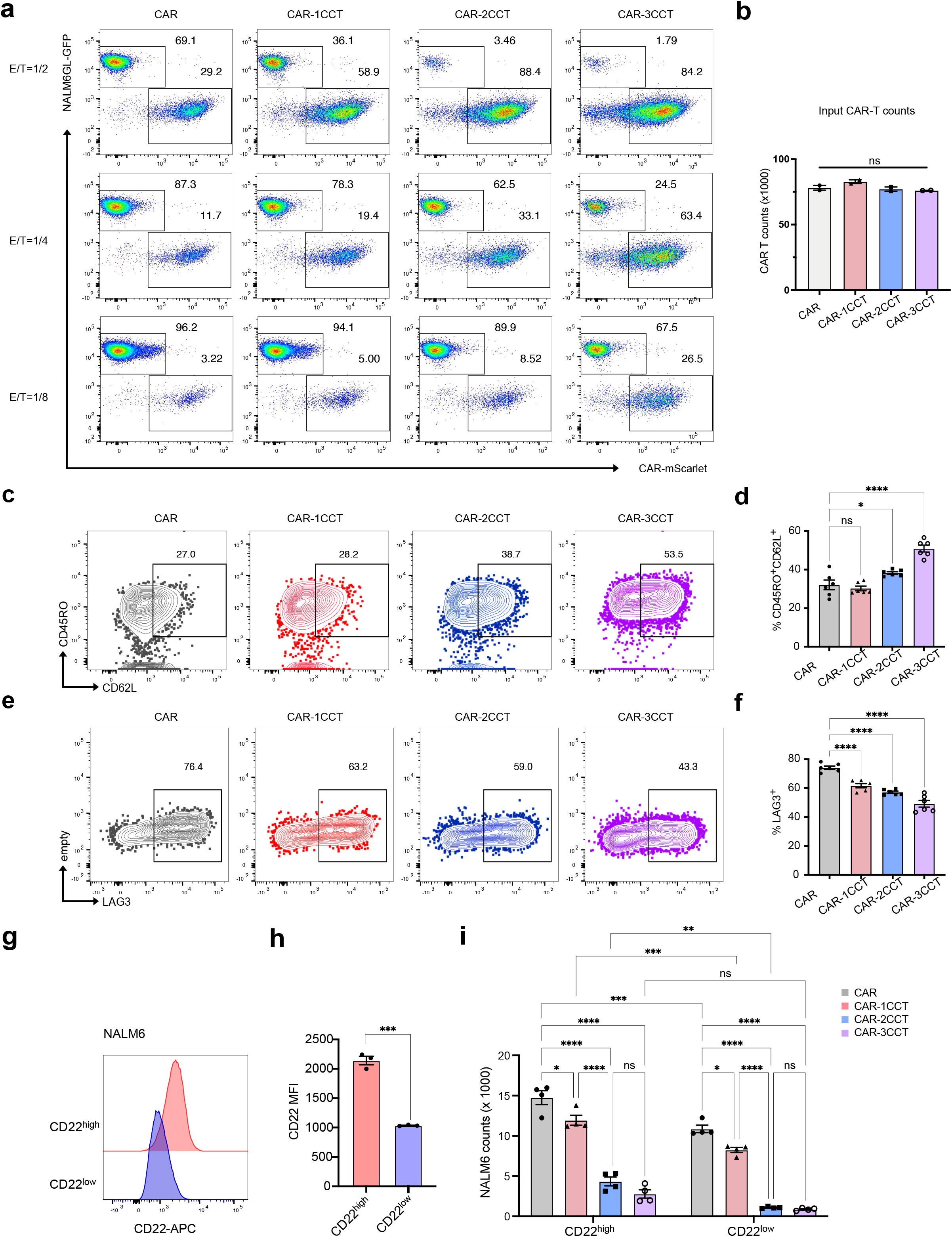
CCT fusion enhanced CAR-T killing under repeated stimulation. **a.** Representative flow cytometry analysis for Fig. 1b**-e** showing CAR-T killing capability under two rounds of NALM6GL stimulation. Purified CAR-T cells were cocultured with NALM6GL cells at variable E/T ratios with variable number of CAR-T cells and fixed amount of NALM6GL cells. NALM6GL cells were re-supplemented at 24hrs after initial coculture. Cells were then collected and stained at 72hrs after initial coculture. **b.** Quantification of input CAR-T cell counts without NALMG6GL stimulation showing relatively equal amount of input CAR-T cells. **c.** Representative flow cytometry results showing the differentiation stage, indicating by the expression of CD45RO and CD62L, of CAR-T cells after two rounds of coculture with NALM6GL cells. **d.** Quantification of **c.** showing the CD45RO^+^ CD62L^+^ CAR-T cells in percentage. **e.** Representative flow cytometry results showing the expression level of LAG3 on CAR-T cells after being stimulated with two rounds of NALM6GL. **f.** Quantification of **e.** showing the LAG3^+^ CAR-T cells in percentage. **g.** Representative flow cytometry analysis on the CD22 expression level of CD22^high^ and CD22^low^ NALM6 cells. **h.** Quantification of **g.** showing CD22 MFI of CD22^high^ and CD22^low^ NALM6 cells. **i.** Quantification of live NALM6 cell count after two-rounds of coculturing with CAR-T cells at E/T = 1/2 by flow cytometry. For all bar plot figures, data are shown as mean ± s.e.m. One-way ANOVA with Dunnett’s multiple-comparisons test is used to assess significance. ns = *p* > 0.05, * = *p* < 0.05, ** = *p* < 0.01, *** = *p* < 0.001, **** = *p* < 0.0001.

**Figure S3.**
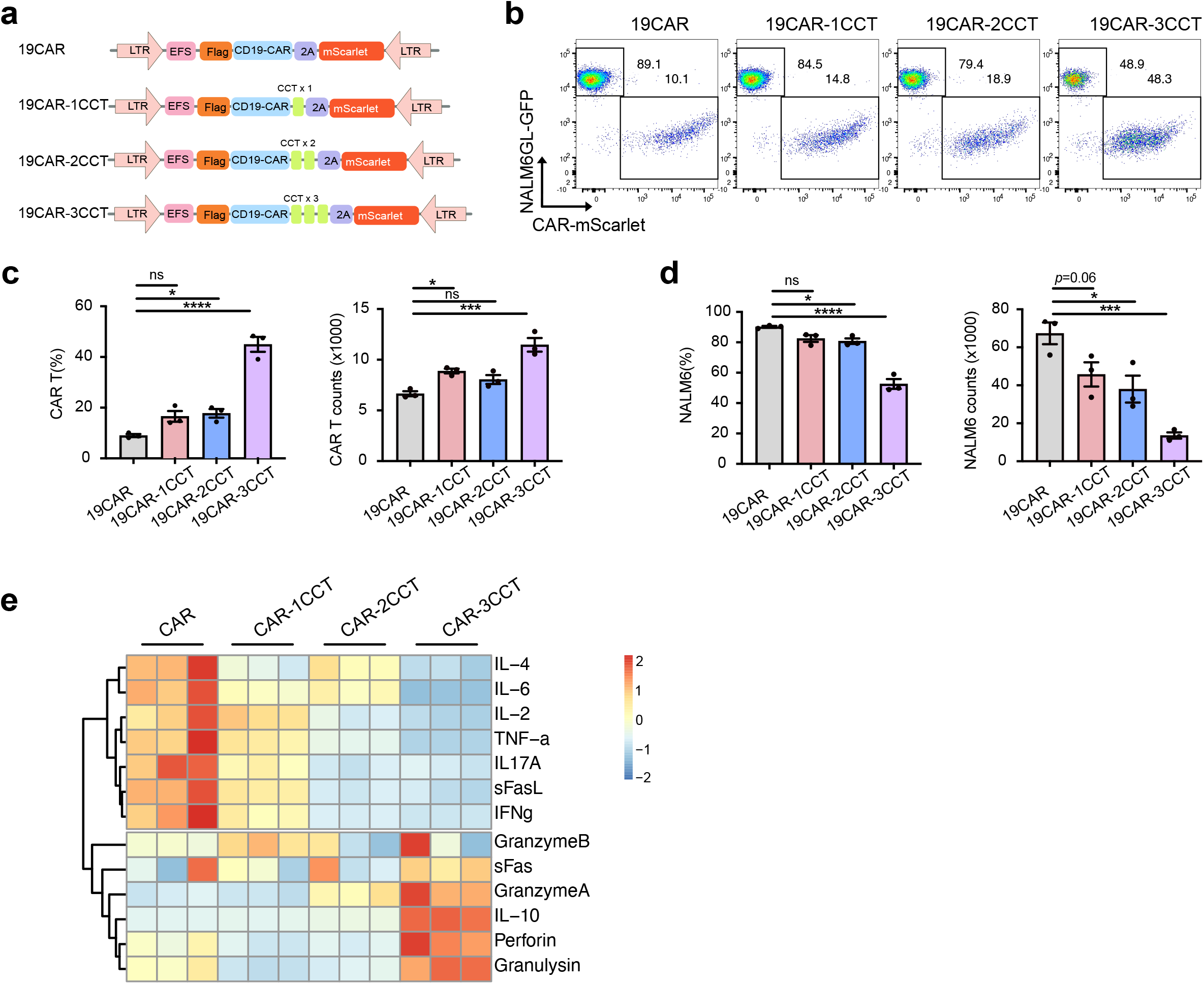
CCT fusion enables enhanced killing with low production of proinflammatory cytokines. **a.** Schematics of lentiviral constructs to make CAR-T cells targeting human CD19 (FMC63-CD28- 41BB-CD3zeta). CD19 CAR (19CAR) constructs were made by replacing the m971 scfv with the FMC63 scgv from Fig. 1a. **b.** Representative flow cytometry analysis of 19CAR-T killing capability *in vitro*. Purified CAR- T cells were cocultured with NALM6GL cells at E/T = 1/4. NALM6GL cells were repeatedly supplemented at 24hrs after initial coculture. Cells were then collected and stained at 48hrs after initial coculture. **c.** Quantification of **b.** showing CD19-CAR percentage (left panel), and counts (right panel). **d.** Quantification of **b.** showing NALM6GL percentage (left panel), and counts (right panel). **e.** Quantification of 13 granule molecules and cytokines in CAR-T and NALM6GL cocultures. CAR-T cells were cocultured with one round of NALM6GL cells at E/T = 1/2 for 24 hours. Coculture supernatant was then collected and measured using LEGENDplex™ Human CD8/NK Panel.

**Figure S4.**
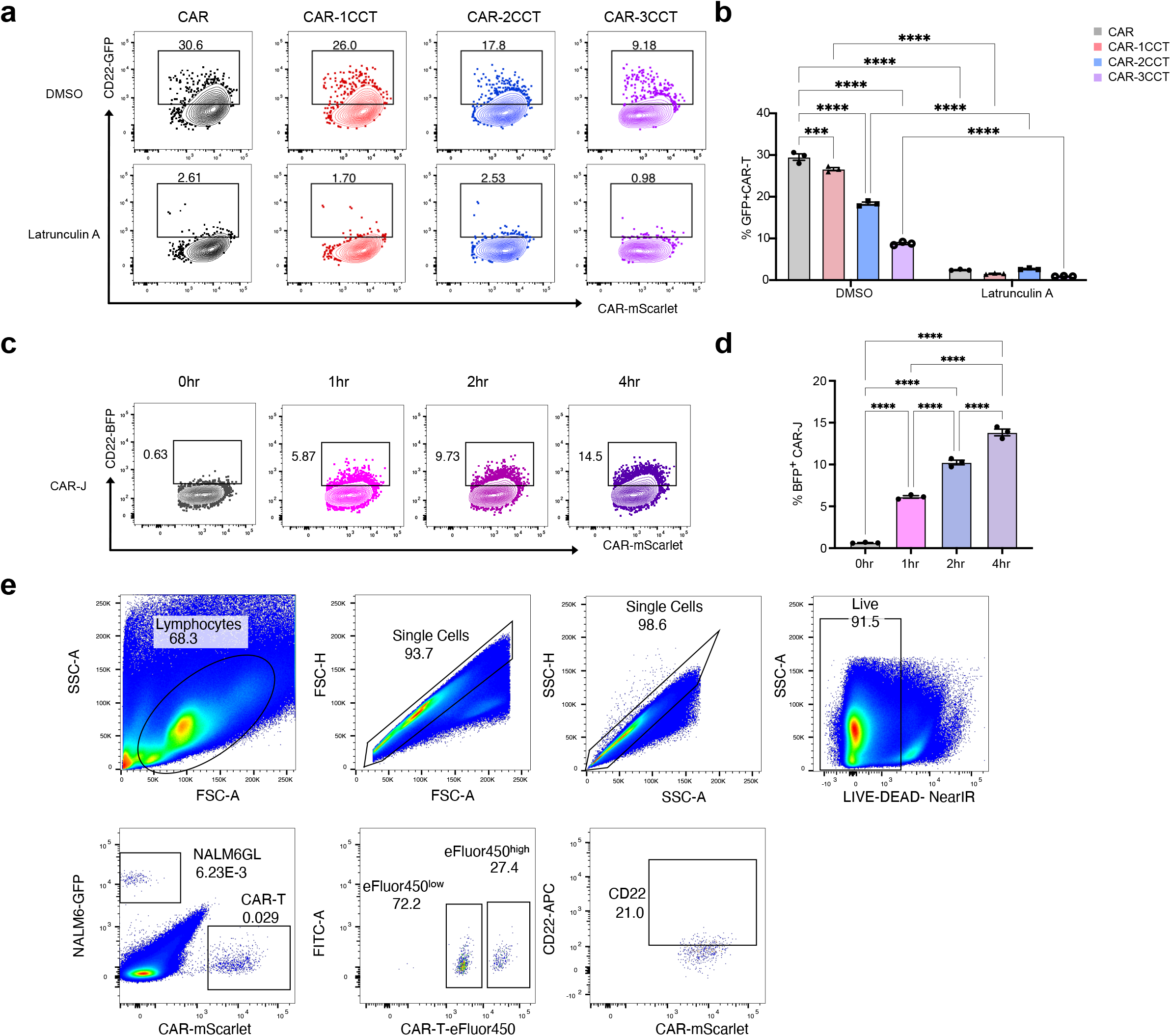
CCT fusion decreased CAR-mediated trogocytosis. **a.** Representative flow cytometry analysis of CAR-mediated trogocytosis. CAR-T cells were pretreated with either DMSO (control) or 1 μM Latrunculin A at 37 °C for 15 min before cocultured with NALM6-CD22-BFP cells for 2 hours. CD22-GFP transfer onto CAR-T cells were quantified by flow cytometry. **b.** Quantification of **a.** showing the inhibition of CD22-GFP transfer by Latrunculin A. **c.** Representative flow cytometry analysis of CAR-mediated trogocytosis by CAR-J cells. CAR-J cells cocultured with NALM6GL-CD22-BFP cells for indicated time. CD22-BFP transfer onto CAR-T cells were quantified by flow cytometry. **d.** Quantification of **c.** showing the transfer of CD22-BFP onto CAR-J cells. **e.** Representative flow cytometry results showing the gating strategy used in Fig. 5 for quantifying the relative CAR-T survival *in vivo*.

**Figure S5.**
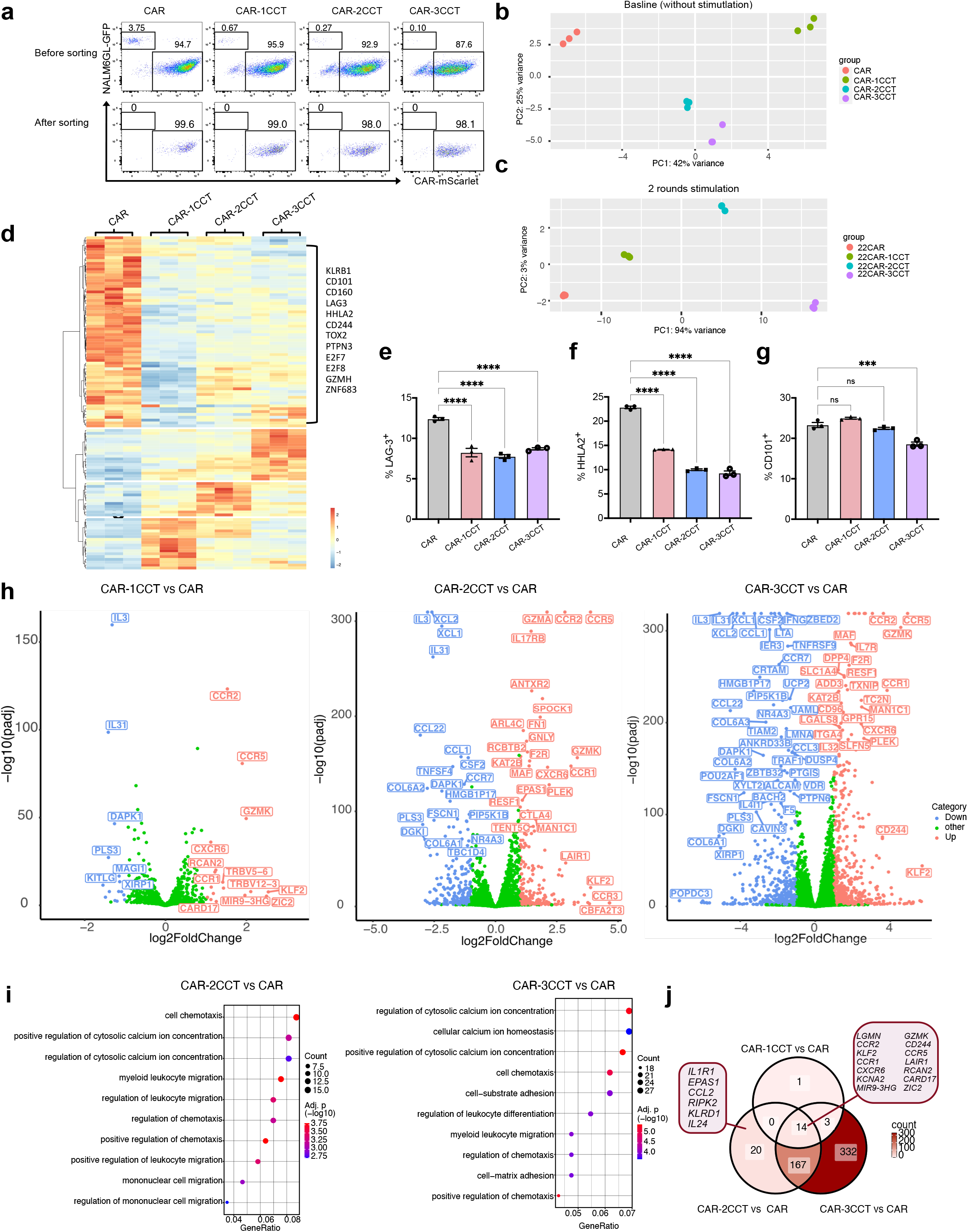
Characterization of CAR-CCT cells by bulk RNA sequencing. **a.** Representative flow cytometry results showing the sorting purity of CAR-T cells before subjecting to RNA-seq. CAR-T cells after being stimulated with 2 rounds of NALM6GL cells were sorted as mScarlet^+^ GFP^-^ cells. **b.** PCA analysis of all four groups of CAR-T cells at baseline without stimulation. **c.** PCA analysis of all four groups of CAR-T cells with 2 rounds of NALM6GL stimulation. **d.** Heatmap of differentially expressed genes in all four groups of CAR-T cells at baseline. All CAR-T cells were purified by flow cytometer at day 6 after lentivirus infection and expanded for 7 days before mRNA extraction for bulk RNA sequencing. Cutoff for determining differentially expressed genes is set to be *p* (adj) < 0.01 and |log2Fc| > 1. **e-g**. Quantification of LAG3(**e**), HHLA2(**f**), CD101(**g**) expression level on CAR-T cells at baseline using flow cytometry. **h.** Volcano plot of bulk RNAseq showing differentially expressed genes in CAR-1CCT vs. CAR (left panel), CAR-2CCT vs. CAR (middle panel) and CAR-3CCT vs. CAR (right panel). Purified CAR-T cells were cocultured with NALM6GL cells at E/T = 1/2. NALM6GL cells were repeatedly supplemented at 24hrs after initial coculture. CAR T cells (mScarlet+) were then sorted at 72hrs after initial coculture for bulk RNAseq. Cutoff for determining differentially expressed genes is set to be *p* (adj) < 0.01 and |log2Fc| > 1. **i.** Pathway analysis by over representation analysis (ORA) showing top 10 pathways of upregulated DEGs in CAR-2CCT vs. CAR (left panel) and in CAR-3CCT vs. CAR (right panel). **j.** Venn diagram showing overlaps of upregulated genes as in (a). Data are shown as mean ± s.e.m. One-way ANOVA with Dunnett’s multiple-comparisons test is used to assess significance. ns = *p*> 0.05, * = *p* < 0.05, ** = *p* < 0.01, *** = *p* < 0.001, **** = *p* < 0.0001.

**Figure S6.**
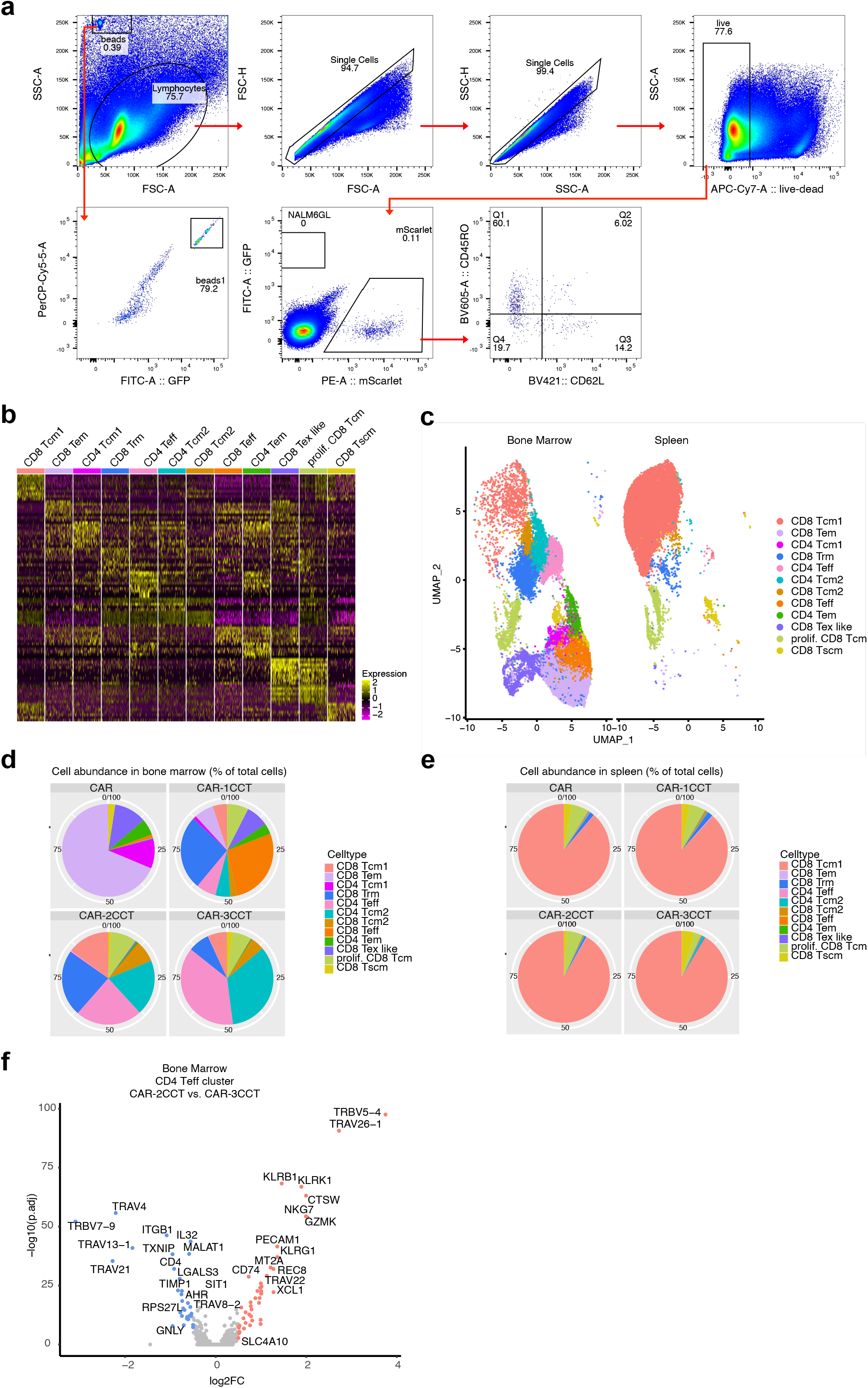
Characterization of CAR-CCT cells *in vivo* via scRNA-seq. **a.** Gating strategy used for analysis shown in Fig. 6 and mScarlet^+^ CAR-T sorting for scRNA-seq. **b.** Heatmap showing expression profiles across 12 CAR-T cell subsets. Scaled gene expression is presented by showing 100 representative cells of each subset. **c.** UMAP visualization of 39,151 cells CAR-T cells via scRNA-seq profiling, split by tissue type. **d.** Pie chart of cell proportions between four different CAR-T groups isolated from bone marrow. **e.** Pie chart of cell proportions between four different CAR-T groups isolated from spleen. **f.** Volcano plot showing differentially expressed genes in CAR-2CCT vs. CAR-3CCT in the CD4 Teff cluster from bone marrow.

**Figure S7.**
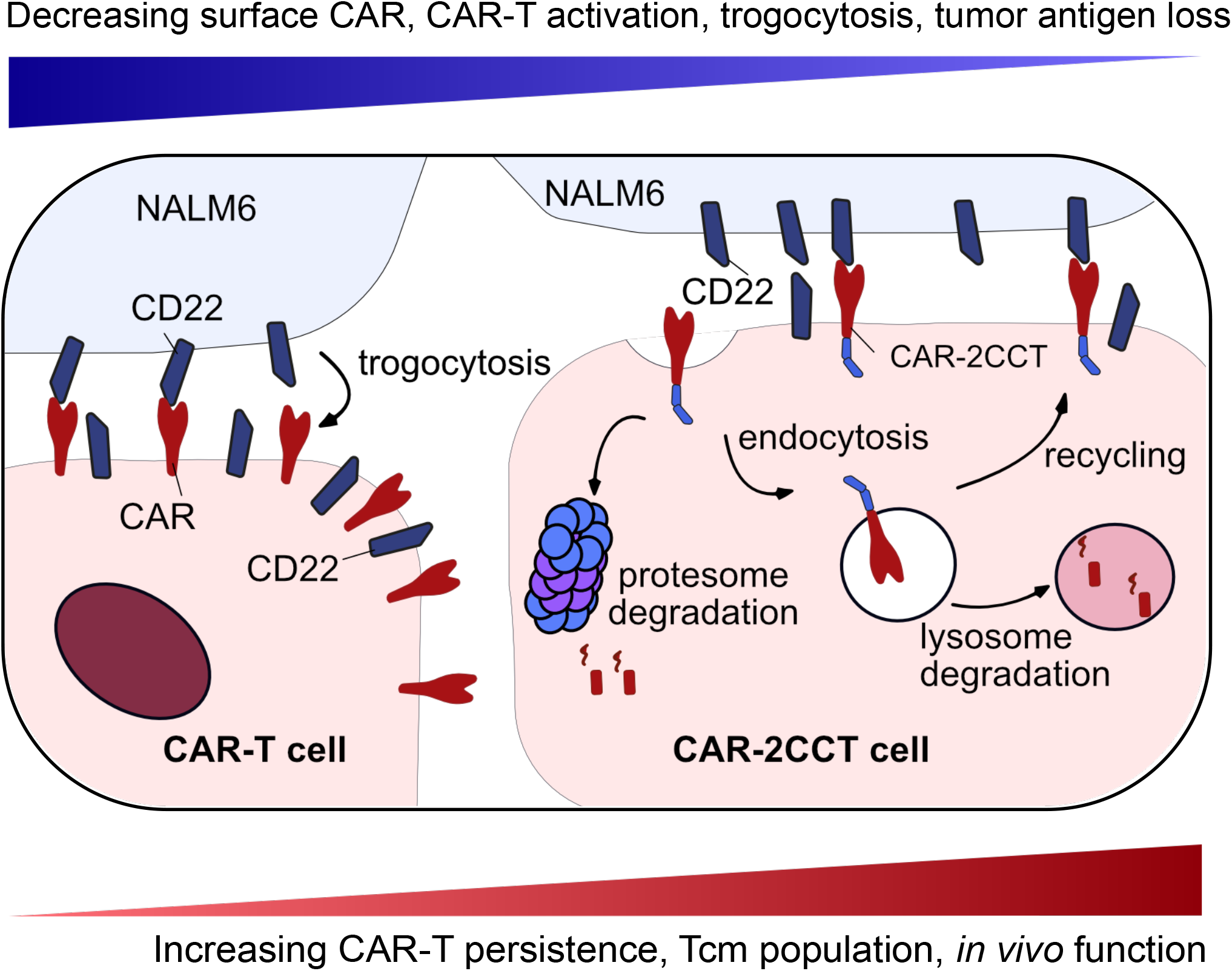
Proposed model of titrating optimal CAR-T function with CCT fusion. CAR fused with duplex CCTs (CAR-2CCT) demonstrated decreased surface expression, which is tightly regulated by its endocytosis, recycling, degradation in both lysosome and proteasome. Compared with the control CAR, CAR-2CCT has significant decreases in surface CAR expression, T cell activation, CAR-mediated trogocytosis, and causes less tumor antigen loss on NALM6 cells. This engineering approach effectively increases the persistence and proportion of Tcm cells in CAR-2CCT cells, leading to a remarkable improvement in their anti-tumor *in vivo*.

## Supplemental Tables and Datasets

### Source data and statistics

Source data and statistics for this study provided in an excel.

## Supplemental Video

**Supplemental Video 1. Time-lapse live cell imaging of trogocytosis and fratricide.** Representative video showing trogocytosis and fratricide of CAR-T cells after introduction with NALM6GL-CD22-BFP cancer cells (Corresponding to Fig. 3a). The white arrowhead labels a CAR-T cell that acquired CD22-BFP after an active engagement with a cancer cell, which subsequently rendered it susceptible to CAR-T fratricide afterwards, as indicated by the influx of SYTOX deep red dye. This white arrow-labeled cell then contacted another CAR-T cell, interacting at a region with enhanced CD22-BFP signal. Shortly after this active interaction, the white arrowhead-labeled cell was stained by the SYTOX deep red dye, indicating cell death. The time stamps on the top right of each image are represented as hour: minute, scale bar = 10µm.

## NGS datasets

**Supplemental Dataset 1. Processed NGS data and analysis results for bulk RNA sequencing.**

Tabs in this dataset:

- Metadata of RNA-seq samples
- Normalized count table by DEseq2 for CAR-T cells at without cancer stimulation (baseline).
- Normalized count table by DEseq2 for CAR-T cells with 2 rounds of NALM6GL stimulation
- Result of differential expression analysis between CAR-1CCT and CAR-T cells at baseline.
- Result of differential expression analysis between CAR-2CCT and CAR-T cells at baseline.
- Result of differential expression analysis between CAR-3CCT and CAR-T cells at baseline.
- Result of differential expression analysis between CAR-1CCT and CAR-T cells after 2 rounds stimulation.
- Result of differential expression analysis between CAR-2CCT and CAR-T cells after 2 rounds stimulation.
- Result of differential expression analysis between CAR-3CCT and CAR-T cells after 2 rounds stimulation.
- List of DEGs after 2 rounds stimulation, q < 0.01 and |log2Fc| > 1 (CAR-1CCT_vs_CAR).
- List of DEGs after 2 rounds stimulation, q < 0.01 and |log2Fc| > 1 (CAR-2CCT_vs_CAR).
- List of DEGs after 2 rounds stimulation, q < 0.01 and |log2Fc| > 1 (CAR-3CCT_vs_CAR).
- Pathway analysis of up-regulated DEGs after 2 rounds stimulation (CAR-2CCT_vs_CAR).
- Pathway analysis of up-regulated DEGs after 2 rounds stimulation (CAR-3CCT_vs_CAR).

**Supplemental Dataset 2. Compiled single cell gene differential expression data.**

Tabs in this dataset:

- Metadata of scRNA-seq dataset.
- Differentially expressed genes across all identified cell types.
- Marker genes and their expression level used for cell type identification.
- Percentages of each cell type in the bone marrow.
- Percentages of each cell type in the spleen.
- List of DEGs showed in the spleen_CD8 Tcm1 (CAR-2CCT_vs_CAR-3CCT).
- List of DEGs showed in the bone marrow_CD4 Tcm2 (CAR-2CCT_vs_CAR-3CCT).
- List of DEGs showed in the bone marrow_CD8 Trm (CAR-2CCT_vs_CAR-3CCT).
- List of DEGs showed in the bone marrow_prolif. CD8 Tcm (CAR-2CCT_vs_CAR-3CCT).
- List of DEGs showed in the bone marrow_prolif. CD4 Teff (CAR-2CCT_vs_CAR-3CCT).

## References

1 June, C. H. & Sadelain, M. Chimeric Antigen Receptor Therapy. N Engl J Med 379, 64–73, doi:10.1056/NEJMra1706169 (2018).

2 Orlando, E. J. et al. Genetic mechanisms of target antigen loss in CAR19 therapy of acute lymphoblastic leukemia. Nat Med 24, 1504–1506, doi:10.1038/s41591-018-0146-z (2018).

3 Majzner, R. G. & Mackall, C. L. Tumor Antigen Escape from CAR T-cell Therapy. Cancer Discov 8, 1219–1226, doi:10.1158/2159-8290.CD-18-0442 (2018).

4 Jayaraman, J. et al. CAR-T design: Elements and their synergistic function. EBioMedicine 58, 102931, doi:10.1016/j.ebiom.2020.102931 (2020).

5 Savoldo, B. et al. CD28 costimulation improves expansion and persistence of chimeric antigen receptor-modified T cells in lymphoma patients. J Clin Invest 121, 1822–1826, doi:10.1172/JCI46110 (2011).

6 Guedan, S. et al. Enhancing CAR T cell persistence through ICOS and 4-1BB costimulation. JCI Insight 3, doi:10.1172/jci.insight.96976 (2018).

7 Weber, E. W. et al. Transient rest restores functionality in exhausted CAR-T cells through epigenetic remodeling. Science 372, doi:10.1126/science.aba1786 (2021).

8 Ghorashian, S. et al. Enhanced CAR T cell expansion and prolonged persistence in pediatric patients with ALL treated with a low-affinity CD19 CAR. Nat Med 25, 1408–1414, doi:10.1038/s41591-019-0549-5 (2019).

9 Daniels, K. G. et al. Decoding CAR T cell phenotype using combinatorial signaling motif libraries and machine learning. Science 378, 1194–1200 (2022).

10 Goodman, D. B. et al. Pooled screening of CAR T cells identifies diverse immune signaling domains for next-generation immunotherapies. Science Translational Medicine 14, eabm1463 (2022).

11 Huang, J. F. et al. TCR-Mediated internalization of peptide-MHC complexes acquired by T cells. Science 286, 952–954, doi:10.1126/science.286.5441.952 (1999).

12 Schriek, P. et al. Marginal zone B cells acquire dendritic cell functions by trogocytosis. Science 375, eabf7470, doi:10.1126/science.abf7470 (2022).

13 Lu, T. et al. Hijacking TYRO3 from Tumor Cells via Trogocytosis Enhances NK-cell Effector Functions and Proliferation. Cancer Immunol Res 9, 1229–1241, doi:10.1158/2326-6066.CIR-20-1014 (2021).

14 Kedl, R. M., Schaefer, B. C., Kappler, J. W. & Marrack, P. T cells down-modulate peptide-MHC complexes on APCs in vivo. Nat Immunol 3, 27–32, doi:10.1038/ni742 (2002).

15 Hamieh, M. et al. CAR T cell trogocytosis and cooperative killing regulate tumour antigen escape. Nature 568, 112–116, doi:10.1038/s41586-019-1054-1 (2019).

16 Olson, M. L. et al. Low-affinity CAR T cells exhibit reduced trogocytosis, preventing fratricide and antigen-negative tumor escape while preserving anti-tumor activity. bioRxiv, 2021.2012.2005.471117, doi:10.1101/2021.12.05.471117 (2021).

17 Rurik, J. G. et al. CAR T cells produced in vivo to treat cardiac injury. Science 375, 91–96, doi:10.1126/science.abm0594 (2022).

18 Tivol, E. A. et al. Loss of CTLA-4 leads to massive lymphoproliferation and fatal multiorgan tissue destruction, revealing a critical negative regulatory role of CTLA-4. Immunity 3, 541–547, doi:10.1016/1074-7613(95)90125-6 (1995).

19 Walunas, T. L. et al. CTLA-4 can function as a negative regulator of T cell activation. Immunity 1, 405–413, doi:10.1016/1074-7613(94)90071-x (1994).

20 Bachmann, M. F., Kohler, G., Ecabert, B., Mak, T. W. & Kopf, M. Cutting edge: lymphoproliferative disease in the absence of CTLA-4 is not T cell autonomous. J Immunol 163, 1128–1131 (1999).

21 Qureshi, O. S. et al. Trans-endocytosis of CD80 and CD86: a molecular basis for the cell-extrinsic function of CTLA-4. Science 332, 600–603, doi:10.1126/science.1202947 (2011).

22 Tekguc, M., Wing, J. B., Osaki, M., Long, J. & Sakaguchi, S. Treg-expressed CTLA-4 depletes CD80/CD86 by trogocytosis, releasing free PD-L1 on antigen-presenting cells. Proc Natl Acad Sci U S A 118, doi:10.1073/pnas.2023739118 (2021).

23 Corse, E. & Allison, J. P. Cutting edge: CTLA-4 on effector T cells inhibits in trans. J Immunol 189, 1123–1127, doi:10.4049/jimmunol.1200695 (2012).

24 Walker, L. S. & Sansom, D. M. The emerging role of CTLA4 as a cell-extrinsic regulator of T cell responses. Nat Rev Immunol 11, 852–863, doi:10.1038/nri3108 (2011).

25 Benmebarek, M. R. et al. Killing Mechanisms of Chimeric Antigen Receptor (CAR) T Cells. Int J Mol Sci 20, doi:10.3390/ijms20061283 (2019).

26 Turtle, C. J. et al. Immunotherapy of non-Hodgkin’s lymphoma with a defined ratio of CD8+ and CD4+ CD19-specific chimeric antigen receptor-modified T cells. Sci Transl Med 8, 355ra116, doi:10.1126/scitranslmed.aaf8621 (2016).

27 Grupp, S. A. et al. Chimeric antigen receptor-modified T cells for acute lymphoid leukemia. N Engl J Med 368, 1509–1518, doi:10.1056/NEJMoa1215134 (2013).

28 Xu, Y. et al. A novel antibody-TCR (AbTCR) platform combines Fab-based antigen recognition with gamma/delta-TCR signaling to facilitate T-cell cytotoxicity with low cytokine release. Cell discovery 4, 62 (2018).

29 Gust, J. et al. Endothelial Activation and Blood–Brain Barrier Disruption in Neurotoxicity after Adoptive Immunotherapy with CD19 CAR-T CellsNeurotoxicity Associated with CD19 CAR-T Cells. Cancer discovery 7, 1404–1419 (2017).

30 Qureshi, O. S. et al. Constitutive clathrin-mediated endocytosis of CTLA-4 persists during T cell activation. Journal of Biological Chemistry 287, 9429–9440 (2012).

31 Sansom, D. M. Moving CTLA-4 from the trash to recycling. Science 349, 377–378 (2015).

32 Janman, D. et al. Regulation of CTLA-4 recycling by LRBA and Rab11. Immunology 164, 106–119 (2021).

33 Kao, S. H. et al. Analysis of Protein Stability by the Cycloheximide Chase Assay. Bio Protoc 5, doi:10.21769/BioProtoc.1374 (2015).

34 Kennedy, A. et al. Differences in CD80 and CD86 transendocytosis reveal CD86 as a key target for CTLA- 4 immune regulation. Nature Immunology 23, 1365–1378 (2022).

35 Martínez-Martín, N. et al. T cell receptor internalization from the immunological synapse is mediated by TC21 and RhoG GTPase-dependent phagocytosis. Immunity 35, 208–222 (2011).

36 Olson, M. L. et al. Low-affinity CAR T cells exhibit reduced trogocytosis, preventing rapid antigen loss, and increasing CAR T cell expansion. Leukemia 36, 1943–1946 (2022).

37 Bloemberg, D. et al. A high-throughput method for characterizing novel chimeric antigen receptors in Jurkat cells. Molecular Therapy-Methods & Clinical Development 16, 238–254 (2020).

38 Huang, F. L., Liao, E. C., Li, C. L., Yen, C. Y. & Yu, S. J. Pathogenesis of pediatric B-cell acute lymphoblastic leukemia: Molecular pathways and disease treatments. Oncol Lett 20, 448–454, doi:10.3892/ol.2020.11583 (2020).

39 Gomes-Silva, D. et al. Tonic 4-1BB Costimulation in Chimeric Antigen Receptors Impedes T Cell Survival and Is Vector-Dependent. Cell Rep 21, 17–26, doi:10.1016/j.celrep.2017.09.015 (2017).

40 Long, A. H. et al. 4-1BB costimulation ameliorates T cell exhaustion induced by tonic signaling of chimeric antigen receptors. Nat Med 21, 581–590, doi:10.1038/nm.3838 (2015).

41 Lee, P. H. et al. Host conditioning with IL-1beta improves the antitumor function of adoptively transferred T cells. J Exp Med 216, 2619–2634, doi:10.1084/jem.20181218 (2019).

42 Velica, P. et al. Modified Hypoxia-Inducible Factor Expression in CD8(+) T Cells Increases Antitumor Efficacy. Cancer Immunol Res 9, 401–414, doi:10.1158/2326-6066.CIR-20-0561 (2021).

43 Jespersen, H. et al. Clinical responses to adoptive T-cell transfer can be modeled in an autologous immune-humanized mouse model. Nat Commun 8, 707, doi:10.1038/s41467-017-00786-z (2017).

44 Maus, M. V. CD19 CAR T cells for adults with relapsed or refractory acute lymphoblastic leukaemia. Lancet 398, 466–467, doi:10.1016/S0140-6736(21)01289-7 (2021).

45 Li, Y. et al. KIR-based inhibitory CARs overcome CAR-NK cell trogocytosis-mediated fratricide and tumor escape. Nature medicine 28, 2133–2144 (2022).

46 Li, G. et al. T cell antigen discovery via trogocytosis. Nature methods 16, 183–190 (2019).

47 Stojanovic, A., Fiegler, N., Brunner-Weinzierl, M. & Cerwenka, A. CTLA-4 is expressed by activated mouse NK cells and inhibits NK Cell IFN-γ production in response to mature dendritic cells. The Journal of Immunology 192, 4184–4191 (2014).

48 Lougaris, V. et al. CTLA-4 regulates human Natural Killer cell effector functions. Clinical immunology (Orlando, Fla.) 194, 43–45 (2018).

49 Philip, M. et al. Chromatin states define tumour-specific T cell dysfunction and reprogramming. Nature 545, 452–456, doi:10.1038/nature22367 (2017).

50 Caruso, H. G. et al. Tuning Sensitivity of CAR to EGFR Density Limits Recognition of Normal Tissue While Maintaining Potent Antitumor Activity. Cancer Res 75, 3505–3518, doi:10.1158/0008-5472.CAN-15-0139 (2015).

51 Rodriguez-Marquez, P. et al. CAR density influences antitumoral efficacy of BCMA CAR T cells and correlates with clinical outcome. Science Advances 8, eabo0514 (2022).

52 Guo, T., Ma, D. & Lu, T. K. Sense-and-respond payload delivery using a novel antigen-inducible promoter improves suboptimal CAR-T activation. ACS Synthetic Biology 11, 1440–1453 (2022).

53 Greenshpan, Y. et al. Synthetic promoters to induce immune-effectors into the tumor microenvironment. Communications Biology 4, 143 (2021).

54 Dai, X. et al. One-step generation of modular CAR-T cells with AAV-Cpf1. Nat Methods 16, 247–254, doi:10.1038/s41592-019-0329-7 (2019).

55 Dobin, A. et al. STAR: ultrafast universal RNA-seq aligner. Bioinformatics 29, 15–21, doi:10.1093/bioinformatics/bts635 (2013).

56 Love, M. I., Huber, W. & Anders, S. Moderated estimation of fold change and dispersion for RNA-seq data with DESeq2. Genome Biol 15, 550, doi:10.1186/s13059-014-0550-8 (2014).

57 Yu, G. enrichplot: Visualization of Functional Enrichment Result. R package version 1.8.1. https://github.com/GuangchuangYu/enrichplot (2020).

58 Satija, R., Farrell, J. A., Gennert, D., Schier, A. F. & Regev, A. Spatial reconstruction of single-cell gene expression data. Nat Biotechnol 33, 495–502, doi:10.1038/nbt.3192 (2015).

59 Jordao, M. J. C. et al. Single-cell profiling identifies myeloid cell subsets with distinct fates during neuroinflammation. Science 363, 365-+, doi:ARTN eaat7554 10.1126/science.aat7554 (2019).

60 Stuart, T. et al. Comprehensive Integration of Single-Cell Data. Cell 177, 1888–1902.e1821, doi:10.1016/j.cell.2019.05.031 (2019).

61 Becht, E. et al. Dimensionality reduction for visualizing single-cell data using UMAP. Nat Biotechnol, doi:10.1038/nbt.4314 (2018).

